# ApTOLL, a new therapeutic aptamer for cytoprotection and (re)myelination after Multiple Sclerosis

**DOI:** 10.1101/2023.01.22.524916

**Authors:** Beatriz Fernández-Gómez, Miguel A. Marchena, David Piñeiro, Paula Gómez-Martín, Estefanía Sánchez, Yolanda Laó, Gloria Valencia, Sonia Nocera, Rocío Benítez-Fernández, Ana M. Castaño-León, Alfonso Lagares, Macarena Hernández-Jiménez, Fernando de Castro

## Abstract

ApTOLL is an aptamer specifically designed to antagonize Toll-Like Receptor 4 (TLR4), a relevant actor for innate immunity involved in inflammatory responses in multiple sclerosis (MS) and other diseases. MS is a primary demyelinating, chronic, inmune and neurodegenerative disease of the central nervous system that normally debuts in young adults. The currently available therapeutic arsenal to treat MS is composed of immunomodulators but, to date, there are no (re)myelinating drugs available in clinics. Our present study shows cells expressing TLR4 in demyelinating lesions of MS patients (*postmortem* samples from cerebral cortex) and, as a derivative, we studied the effect of TLR4 inhibition with ApTOLL in animal models of MS (experimental autoimmune encephalomyelitis -EAE- and the cuprizone). The treatment with ApTOLL positively impacted the clinical symptomatology, and this was associated with better preservation plus restoration of myelin and oligodendrocytes in the demyelinated lesions of these animals, which suggests not only an immunomodulatory but also a remyelinating effect of the treatment with ApTOLL. This latter was corroborated on purified cultures of rodent and adult human oligodendrocyte precursor cells (OPCs), confirming the expression of TLR4 in this cell type. Altogether, the molecular nature of ApTOLL and its mechanism/s of action strongly supports this compound as a novel candidate to treat MS and other demyelinating scenarios.

## INTRODUCTION

Multiple sclerosis (MS) is a primary demyelinating, chronic, neurodegenerative, inflammatory and autoimmune disease of the central nervous system (CNS). It represents the leading cause of non-traumatic disability in young adults, first displaying clinically signs around the age of 20-40 years [59]. As in other primary demyelinating diseases, the main pathophysiological feature of MS is the loss of oligodendrocytes and myelin in the CNS, both in the white (WM) and gray matter (GM) [77]. This process is also associated with the activation of macrophages and microglia, the resident innate immune cells in the CNS, causing a background of inflammatory reactions [2, 40]. After damage, endogenous OPCs spontaneously migrate to lesion sites, where they can differentiate into new myelin-forming mature oligodendrocytes [13, 19, 74]. Although it is known that microglia can promote remyelination through the clearance of myelin debris, the secretion of growth factors and cytokines, and the modulation of the extracellular matrix, chronic activation of microglia contributes to the progression of neurodegeneration through the release of multiple neurotoxic factors like tumor necrosis factor-α, nitric oxide or reactive oxygen species [4, 44]. The currently available treatments to manage MS are based on the prescription of immunosuppressive and immunomodulatory agents, although there is still no approved compound to successfully induce disease-reversing remyelination [23, 39]. The search for remyelinating approaches is a hot challenge for modern Neurology, particularly since the first clinical trial showed positive effects of such an approach [24]. The experimental autoimmune encephalomyelitis ( E A E ) i s t h e m o s t frequently used animal model to study MS since it induces a variety of immunopathological and neuropathological events that approximate closely to the key features of the relapsing-remitting variety of MS (RRMS) [10], while in the cuprizone model it is possible to study demyelination and remyelination without the intervention of the immune system [25].

The family of Toll-Like Receptors (TLRs) participates in innate immunity, stimulating a wide variety of inflammatory responses [52]. Specifically, TLR4 has been implicated in several pathologies with a significant inflammatory component, including MS [7, 37, 84]. TLR4 activation increases the expression and nuclear translocation of nuclear transcription factor kappa-B (NF-kB), leading to the release of proinflammatory cytokines like TNF-α, interleukin-1β (IL-1β) or interleukin-6 (IL-6), as well as chemokine and lymphocyte recruitment [43, 45]. In this regard, TLR4 inhibition is known to enhance the survival of oligodendrocytes and OPCs [29, 41, 76]. Therefore, the possibility of down-regulating immune responses with specific TLR antagonists, inhibiting specific intracellular proteins involved in these signaling pathways, has raised great interest.

This is the case of ApTOLL, a specific innovative antagonist of TLR4 with the potential to achieve an immunomodulatory and anti-inflammatory effect. ApTOLL is a single-stranded DNA aptamer selected using the systemic evolution of ligands by exponential enrichment (SELEX) technology, and its molecular nature and mechanism of action offers a novel strategy to treat diseases with a significant inflammatory component, substantially reducing brain damage [16]. Aptamers offer several advantages over antibodies that make them ideal for future therapeutic applications, such as their high specificity and affinity, and their ease to enter biological compartments given their small size and lack of immunogenicity [86]. Therefore, based on its pharmacokinetic characteristics and its very low level of toxicity, ApTOLL would be suitable for the treatment of different diseases. In fact, ApTOLL has been successfully tested in preclinical models of diseases like ischemic stroke and myocardial infarction, producing an excellent protective effect [16, 63, 69]. In clinical studies, ApTOLL has demonstrated a very good safety profile in a completed First-in-Human clinical trial [31] and it is currently undergoing a Phase Ib/IIa trial (APRIL trial) to determine its safety and biological effect in acute ischemic stroke patients [30].

In the present work, we are the first to demonstrate cells expressing TLR4 in human demyelinating lesions from MS patients, as well as the beneficial effect of this aptamer in EAE and cuprizone models of MS. These results *in vivo* were completed with a detailed *in vitro* study of the effects of ApTOLL on primary cultures of OPCs, as well on the myelination in organotypic cultures of cerebellar slices. Together, our present proof-of-concept candidates ApTOLL as a therapeutic agent to treat MS, possibly in combination therapy to obtain immunomodulatory, neuroprotective and neuroreparative effects. In such a scenario, ApTOLL would reduce inflammation and demyelination, as well as promote the survival and replenishment of endogenous OPCs and mature oligodendrocytes, thereby enhancing remyelination.

## MATERIALS AND METHODS

### Human brain samples

*Post-mortem* cortical brain tissue blocks from human MS patients and controls were obtained from the UK Multiple Sclerosis Tissue Bank. Alternate coronal slices or brain hemispheres were fixed in 4% paraformaldehyde (PFA) for∼2 weeks and were then cryoprotected in 30% sucrose for ∼1 week and frozen in isopentane precooled on a bed of dry ice. Microtome sections (50 μm) were obtained from all the cortical blocks containing subjacent WM for immunohistochemical analysis [8]. Blocks from MS patients with WM lesions and controls were studied here, from individuals with no history of neuropsychiatric disease in either case (Table 1). These experiments with human samples have been approved by the CSIĆs Comité de Ética (073/2021).

### Animals

Female and male C57/BL6 mice (7-weeks-old) were purchased from Charles River Laboratories (Wilmington, MA, USA), while C57BL/10ScNJ mice (formerly C57BL/10ScCr) that do not express TLR4 (TLR4^-/-^) [66] were purchased from Jackson Laboratory (Bar Harbor, Me, USA) and the colony was maintained at the *Centro de Investigaciones Biológicas Margarita Salas (CIB)-CSIC* . The animals used in the *in vivo* studies were transferred to the animal facility at the *Instituto Cajal*, where the mice were acclimatized for a week in the appropriate environmental conditions with *ad libitum* access to food and water prior to carry out the procedures. Wistar P7 rats from the animal facility of the *Instituto Cajal* were also used in primary cultures. All experimental procedures were performed in accordance with European Council Guidelines (63/2010/EU), and Spanish National and Regional Guidelines for Animal Experimentation and Use of Genetically Modified Organisms (RD 53/2013 and 178/2004, Ley 32/2007 and 9/2003, Decree 320/2010). The generation of the different MS models was approved by the relevant institutional and regional ethics committees CSIC (440/2016, Madrid, Spain).

### Induction of active EAE

EAE was induced in female *wild type* (WT) mice as described previously [53, 56]. The mice were first anesthetized with 40 µl of an anesthetic/analgesic solution containing Ketamine (40 mg/ml: Anesketin) and Xylazine (2 mg/ml: Rompun) injected intraperitoneally (i.p.). An emulsion of the Myelin Oligodendrocyte Glycoprotein (MOG_35-55_ peptide, 250 µg in a final volume of 200 µl: GenScript) and complete Freund’s adjuvant (CFA) containing inactivated *Mycobacterium tuberculosis* (4 mg: BD Biosciences) was introduced subcutaneously into the animal’s groin and armpits. Then, pertussis toxin (400 ng/mouse: Sigma-Aldrich) was administered intravenously (i.v.) through the tail vein on the day of immunization and 48h later. The mice were weighed and evaluated double-blind on a daily basis until sacrifice, scoring the animals as follows: 0, asymptomatic; 1, loss of muscle tone throughout the tail; 2, weakness or unilateral partial hind limb paralysis; 3, bilateral paralysis of the hind limbs; 4, tetraplegia; and 5, death. In all cases, mice were sacrificed ten days after the onset of symptoms. After EAE induction, animals were evaluated on a daily basis and sacrificed 10 days after the onset of clinical symptoms.

### ApTOLL treatment

ApTOLL (0.45 mg/kg, 0.91 mg/kg, 1.82 mg/kg or 3.6 mg/kg, diluted in sterile PBS/1mM MgCl _2_ -vehicle) was administered i.v. to the WT and TLR4^-/-^ mice through a single injection at the onset of the symptoms (defined as a clinical score between 0.5 and 1.5). For the study of the therapeutic window, independent groups were established, and a single 0.91 mg/Kg dose was injected 24h after the onset and at the peak of the symptoms.

### Cuprizone-induced model

Male, 8-week-old C57/BL6 mice were randomly separated into three groups: control untreated, ApTOLL-treated, and vehicle treated. While the control animals received a normal diet, the rest of the mice were fed chow containing cuprizone (CPZ, bis (cyclohexanone) oxaldihydrazone: Sigma-Aldrich) at 0.25% (w/w) *ad libitum* for 6 weeks [51], a copper chelator that induces apoptosis of mature oligodendrocytes and that leads to robust demyelination, particularly in the corpus callosum (CC) [25, 67]. The body weight of each mouse was controlled weekly, and ApTOLL or the vehicle alone was also injected i.v. once weekly from the beginning of the experiments to the time of sacrifice. To study CPZ-induced oligodendroglial proliferation, we add EdU (5-ethynyl-2’-deoxyuridine; Invitrogen), a direct measure of de novo DNA synthesis, into the drinking water at 0.2 mg/ml the week before sacrifice.

### Motor coordination test (Rotarod)

At the end of the 6-week experimental period, the animals were tested on the rotarod to evaluate their locomotor coordination [17]. For training, mice were subjected to pre-test trials the day before, placing them on the rod at constant speed (30 rpm). Each mouse performed the test twice in the same conditions as the pre-test trials and the latency to fall was measured.

### Tissue sampling

Animals were sacrificed by i.p. administration of a lethal dose of pentobarbital and they were perfused transcardially with 4% PFA in 0.1M Phosphate Buffer (PB, pH 7.4). The brain and spinal cord were obtained and post-fixed for 4h at room temperature (RT) in 4% PFA. The tissue was cryoprotected by immersion in an increasing sucrose gradient and coronal cryostat sections (20 μm) were obtained on a CM1900 cryostat (Leica).

### Eriochrome-cyanine staining

To analyze CNS demyelination, eriochrome-cyanine (EC) staining was performed as described previously [51, 56]. The histological sections were dried for 2h at RT and for 2h at 37 °C, before they were immersed in acetone for 5 min, dried and submerged in the EC solution (Sigma) for 30 min. To differentiate the samples, iron alum at 5% (w/v) was used for 5– 10 min followed by a second differentiation step with borax-ferricyanide solution for 5 min at RT. For conservation, the sections were dehydrated in ethanol solutions of increasing concentration and mounted with mounting medium (DPX, Sigma).

### Immunohistochemistry in CNS tissue

In human tissue, TLR4 and HLA-DR was detected using diaminobenzidine (DAB). Briefly, the tissue was first pre-treated with 0.3% hydrogen peroxide in methanol for 5 min in order to eliminate endogenous peroxidases. Samples were washed in 0.1M PB and blocked with a solution containing: 1% of normal goat serum (NGS), 0.2% of Triton X-100 in PBS for 1h at RT. After that time, they were incubated with the primary antibody (see Table 2) overnight at RT, washed with PBS and incubated with the secondary antibody (1:1000) for 1:30h at RT. Samples were then washed with PBS and immersed in the solution containing the avidin-biotin-peroxidase complex (Ultra-Sensitive ABC Peroxidase Standard Staining Kit, ThermoFisher) for 30 min at RT. Finally, they were washed with PBS and 0.1% Tris-HCl buffer for 5 min, and peroxidase was revealed with a solution consisting of: 0.02% of 3,3’-diaminobenzidine (Sigma), 0.003% of hydrogen peroxide in 0.2% Tris-HCl buffer. The reaction was controlled under the microscope and stopped by washing the samples with 0.05% of Tris-HCl buffer. Before mounting, the tissue was dehydrated in ethanol solutions of increasing concentration.

To detect antigens present in the spinal cord (SC) of EAE mice and the CC from CPZ-treated animals, sections were first pre-treated with 10% methanol in 0.1M PB at RT for 15 min and after several washes with 0.1M PB and PB saline (PBS), then they were incubated for 1h at RT with blocking solution to avoid non-specific binding: 5% of normal donkey serum (NDS) and 0.02% of Triton X-100 (Sigma-Aldrich) in PBS. The sections were then incubated overnight at 4 °C with the primary antibodies (see Table 2) and after washing, they were probed for 1h at RT with the fluorescent secondary antibodies (1:1000: ThermoFisher). Cell nuclei were stained with Hoechst 33342 (10 μg/mL: Sigma-Aldrich) and finally, the tissue was washed and mounted with Fluoromont® mounting medium (SouthernBiotech). We also used FluoroMyelin Green fluorescent myelin stain (Invitrogen) for the quick and selective labeling of myelin in brain cryosections. As such, the samples were rehydrated with PBS and incubated with the staining solution for 20 min at RT. Subsequently, the sections were rinsed in PBS three times and mounted with Fluoromont®. To visualize cell proliferation, EdU detection cocktail (Click-iT EdU Alexa Fluor 488 HCS assay, Invitrogen) was used according to the manufacturer’s instructions.

### Primary Oligodendrocytes precursor cell (OPC) cultures

OPCs from the cerebral cortex of P7 Wistar rats and TLR4 ^-/-^ mice were isolated using Magnetic-Activated Cell Sorting-MACS (Miltenyi) [14, 54]. First, the animal’s brain was extracted and the meninges removed in a Petri dish containing cooled HBSS ^+/+^ (Hank’s Balanced Salt Solution with Ca ^2+^ and Mg ^2+^: Gibco) to maintain their metabolism. According to the manufacturer’s instructions, the brain cortices were then mechanically triturated in HBSS^-/-^ and enzymatically digested for 25 min at 37 °C with the enzymes provided in the Neural Tissue Dissociation Kit (P) (Miltenyi) under mild and continuous agitation. After adding the second mix, the tissue was dissociated with a 5 ml pipette. The suspension was passed through 70 µm nylon cell strainer (BD Falcon), centrifuged at 300 g for 10 min and incubated for 25 min at 4°C with the primary antibody from A2B5 (Millipore) and/or O4 hybridoma cells (DSHB: Developmental Studies Hybridoma Bank, Iowa) diluted 1:1 in a Miltenyi wash buffer (MWB – 2 mM sodium pyruvate, 0.5% BSA and 2 mM EDTA adjusted to pH 7.3). After that, cells were washed with MWB and exposed to the secondary rat anti-mouse IgM antibody (anti-mouse IgM microbeads: Miltenyi) for 151min at 201μl/10^7^ cells. Next, the cell suspensions were washed and passed through a MWB-coated medium-size column (MS, Miltenyi) coupled to a magnet and eluted with Neuromedium (Miltenyi) supplemented with 2% MACS® NeuroBrew-21 w/o vitamin A (Miltenyi), 0.6% Penicillin/Streptomycin (10,000 U/ml penicillin and 10 mg/ml streptomycin: Sigma), glutamine (200 mM, Gibco), D-glucose (25 mM, Normapur) and 10 ng/mL of PDGF-AA (Platelet Derived Growth Factor-AA: Merck) and FGF-2 (Fibroblast Growth Factor-2: Peprotech). Finally, the cells were counted and seeded on coverslips coated with poly-L-Lysine (0.1 mg/ml in borate buffer, pH 8.5: Sigma) and laminin (10 µg/ml in PBS: Sigma). A purity of ∼90% was obtained.

Human OPCs were isolated from biopsies of the adult cerebral cortex following surgery of traumatic brain injury patients, obtained from the Neurosurgery Service of the Hospital 12 de Octubre (Madrid), and conserved immediately in Hibernate-A (Gibco) at 4 °C up to the isolation process. The study was conducted according to the guidelines and the Research Ethics Committee approved protocol of the *Instituto Cajal-CSIC* (440/2016 and 2016/049/CEI3/20160411) and *CSIC* (073/2021). To obtain the cells, the same procedure was applied with some modifications. After filtration, the pellet was passed through a 20% Percoll (GE Healthcare) gradient and centrifuged at 800 g for 20 min. The cells were then exposed to a Red Blood Cell Removal Solution (Miltenyi) diluted in ddH _2_O for 10 min at 4 °C. An anti-O4 Microbead antibody (2.5 μl: Miltenyi) was resuspended in 97.5 μl of MWB and used to label up to 10 ^7^ total cells by incubating for 15 min at 4 °C in the dark with mild agitation. Culture purity was at least 85%.

#### Viability/survival assay

The viability of the OPCs after the different treatments was tested with a MTT (Thiazolyl blue tetrazolium bromide) assay that measures the mitochondrial activity of the cells by quantifying the conversion of the tetrazolium salt to its formazan product. Cells were seeded in 96-well plates at a density of 15,000 cells/well in a final volume of 200 µl and treated the day after the culture with ApTOLL (20 nM and 200 nM), the vehicle alone or 5% H _2_O_2_ as a cell death control. After 24h, MTT (5 mg/ml: Sigma) was added to the plates and the cells were left in the incubator for 4h, after which the medium was removed, DMSO (dimethyl sulfoxide; Invitrogen) was added to dissolve the formazan formed and the absorbance of this solution was measured. Likewise, the survival of OPCs was determined by TUNEL staining (Terminal deoxynucleotidyl transferase dUTP nick end labeling: see [54]. To this end, the cells were seeded in 24-well plates at 30,000 cells/coverslip in a final volume of 500 µl, treated with ApTOLL or the vehicle alone on the following day, and then fixed in 4% PFA after 48h.

#### Proliferation assay

To analyze OPC proliferation, the cells were seeded, as previously described, at a density of 30,000 cells/coverslip [50, 54] and after 24h, the medium was replaced with medium containing the corresponding treatments and 6h a pulse of BrdU (5-bromo-2-deoxyuridine, 50 µM; Sigma-Aldrich) was given to the neonatal cells. At the end of this time, the medium was refreshed and the cells were fixed with 4% PFA on the following day.

#### Differentiation assay

Cells were seeded at 30,000 cells/coverslip and 24h later, the medium was replaced with new medium without growth factors (PDGF-AA and FGF-2). The following day the treatments were added and the cells were fixed and immunolabelled at the end of the fifth day *in vitro*.

In experiments in which inflammatory conditions were induced, IL1-β (50 ng/ul) and TNF-α (50 ng/ul) were added to the culture medium the day after cell isolation and fixed 24h later to determine the TLR4 expression.

### *In vitro* pharmacology

Human TLR2,-3-4-5-7- 8-and 9 expressing HEK293 cell line were incubated with ApTOLL (20nM and 200nM) and their corresponding agonists (PAM2, Poly I:C, LPS K12, Flagellin, R848, R848 and OND2006, respectively) in a specific study performed by Invivogen. These cell lines functionally overexpress a given TLR protein as well as a reporter gene which is a secreted alkaline phosphatase. The production of this reporter gene is driven by a NFkB inducible promoter. A recombinant HEK-293 cell line for the reporter gene only was used as a negative control for the TLR reporter cell lines. This negative control cell line does not express any of these receptor genes, but the reporter gene (alkaline phosphatase) only. This reporter gene is directly inducible with TNFα.

Moreover, SafetyScreen44™ (EUROFINS) panel was performed to enable early identification of significant off-target interactions with ApTOLL. Forty-four targets (GPCRs, Ion Channels, Kinases, Nuclear Receptors, Transporters and other Non-Kinase Enzymes) were selected to bring together both robustness and the strategic choice of information-rich targets. Compound binding was calculated as a % inhibition of the binding of a radioactively labeled ligand specific for each target. Compound enzyme inhibition effect was calculated as a % inhibition of control enzyme activity. In each experiment and, if applicable, the respective reference compound was tested concurrently with ApTOLL, and the data were compared with historical values determined at EUROFINS.

### Immunocytochemistry of *in vitro* cultures

After fixing the cells with 4% PFA, they were washed with PBS and maintained for 1h under shaking in the blocking solution (5% NDS and 0.02% Triton X-100). The cells were then incubated overnight with the corresponding primary antibodies in a humid chamber at 4 °C (Table 2) and after washing they were exposed to the secondary antibodies for 1h at RT in the dark. The nuclei were stained with Hoechst (1:20) for 10 min in the dark and the coverslips were mounted in Immu-mount (ThermoFisher).

To study proliferation, BrdU labeling was performed after Olig2 staining. After washing the cells, they were denatured with HCl 2N for 45 min at RT, washed with borate buffer and blocked for 3h at RT with PBS containing 5% FBS (Fetal Bovine Serum), 0.03% Triton X-100, 0.2% gelatin and 0.2M glycine. They were then incubated overnight at 4 °C with an anti-BrdU rat antibody (Abcam) diluted in blocking solution and after several washes, they were exposed to the secondary antibody for 4h before Hoechst staining. TUNEL staining was performed with the ApoTag Plus Fluorescein *in Situ* Apoptosis kit (Millipore Merck, Sigma-Aldrich) following the manufacturer’s instructions. Briefly, cells were equilibrated with the corresponding buffer and incubated for 1h at 37 °C in 50% Terminal deoxynucleotidyl transferase (Tdt) in reaction buffer. The reaction was stopped and after washing, the coverslips were exposed for 30 min to 47% anti-digoxigenin diluted in the blocking buffer. Finally, the standard protocol was applied to label additional markers.

### Organotypic cultures of cerebellar slices

Cerebellar slices from neonatal mice were cultured following standard protocols [3, 51, 54]. Mice were decapitated, and their brain and cerebellum were deposited in a Petri dish with HBSS^+/+^ on ice. The cerebellum was separated, immediately placed on a Teflon base and cut automatically into 350 μm slices on a Mcllwain Tissue Chopper (Mickle Laboratory Engineering Co. Ltd). Cerebellar parasagittal slices were transferred under sterile conditions to 30 mm diameter MilliCell inserts with a pore size of 0.4 μm (Millipore), placed in a 6-well plate with 1 ml of culture medium consisting of: 25% HBSS ^+/+^, 50% BME (Basal Medium Eagle: Gibco), 25% inactivated horse serum (Gibco), 28 mM D-glucose, 1% Penicillin/Streptomycin solution and 0.25 mM Glutamax (Gibco). The medium was refreshed every other day and a demyelinated toxin, lysolecithin (LPC, 0.5 mg/ml: Sigma), was added after 7 days *in vitro* (DIV) for 15h. Subsequently, the medium was removed and the slices were treated with ApTOLL (20 nM) or the vehicle alone in fresh culture medium. The cultures were maintained until 6 days post-lesion (dpl), refreshing the medium and treatment every other day, fixing samples with 4% PFA for 15 min at both 0 dpl as an internal control for the effectiveness of LPC and at 6 dpl. Finally, they were stained for MBP (Myelin Basic Protein), NFH (Neurofilament heavy chain) and Iba1 as described above.

### Image and data analysis

Optical images of EC and DAB staining were acquired and mosaics built using the brightfield camera of the THUNDER microscope (LEICA) at 20X and 40X magnification. Fluorescence images of tissue samples were taken on an inverted Leica SP-5 confocal microscope at the Microscopy and Image Analysis Unit of the *Instituto Cajal* . In the EAE model, mosaics of three SC samples were obtained per animal at 40x magnification and at a resolution of 512 x 512 pixels, with a separation between z-planes of 3 µm. Demyelinated areas and MBP/NFH staining were assessed using the ImageJ application, and the cell counts analyzed with the microscopy software for 3D and 4D Imaging (IMARIS). Similarly, the central zone of the CC was evaluated in the CPZ model. Images were acquired with the same confocal microscope at a 40x magnification (for Caspr labeling a 63x objective was used), at a resolution of 1024 x 1024 pixels and with a separation of 0.5 µm between the z-planes. The same parameters were applied for microphotographs of cerebellar slices, including a digital zoom of 1.7. The quantification of the different markers was performed using ImageJ software. Images of primary cultures were obtained with a Leica AF 6500-7000 microscope (five random photos per coverslip at 20x magnification) and they were counted manually with the LAS X Life Science software. The absorbance for the viability assay was measured with a fluorometer FLUOstar OPTIMA (BMG Labtech) at 595 nm.

### Morphometric analysis of microglia

We studied the morphological complexity of microglia to determine their functionality in organotypic slices under inflammatory conditions (exposed to LPC). A minimum of 10 cells per condition were randomly analyzed and processed with the ImageJ software, following the previously published protocol [80]. In summary, for the skeleton analysis the image was first adjusted for brightness/contrast if necessary and processed to remove background noise. It was then converted to a binary image and the AnalyzeSkeleton (2D/3D) plugin was run. The results were obtained from the Branch Information box.

### Statistical analysis

The data are expressed as the means ± SEM throughout and they were analyzed with GraphPad (GraphPad Software, La Jolla California USA). First, a D’Agostino & Pearson normality test was passed to check if the values followed a Gaussian distribution. To compare pairs of independent groups, Student’s *t*-test was used or a Mann-Whitney U test for non-parametric data. For multiple comparisons, One-Way ANOVA test was carried out in conjunction with the corresponding *post hoc* tests: Tukey’s test for parametric samples and Dunn’s test for non-parametric samples. The threshold for statistical significance was set at p< 0.05 and the results of the analysis were represented in text and figures as: *p <0.05; ** p <0.01; ***p <0.001.

## RESULTS

### Presence of TLR4^+^ cells in demyelinating plaques of MS patients

TLR4 has been associated with pathological processes and inflammation, hence we studied whether TLR4 was dysregulated in the CNS of *post-mortem* samples of MS patients (Table 1). As in a previous study [8, 70], we identified demyelinating lesions by EC staining in the WM of each of the MS patients studied (Fig. 1B-E), while none was observed in the blocks available from the five control individuals (Fig. 1A). To detect the presence of inflammatory infiltrate, HLA-DR allowed us to identify the different types of lesion, thus distinguishing active (Fig. 2B-D) and chronic-active lesions (Fig. 2E) [8]. As expected, HLA-DR staining was absent in samples from controls patients (Fig. 2A). While almost any track of TLR4 staining was detected in the tissue of control patients (Fig. 3A), TLR4 ^+^ cells were abundant within active lesions (Fig. 3B-D), but just scarce in the core of chronic plaques (Fig. 3E) of MS patients. According to previous studies in similar samples [8, 70], the morphological aspect and distribution of TLR4^+^-cells in active lesions suggested a net inflammatory predominance (Fig. 3B-D), but in the core of chronic lesions, the very few TLR4 ^+^-cells showed different morphologies, suggesting a different cellular nature (Fig. 3E). As TLR4 is upregulated in patients with MS [1, 72], we next investigated this in preclinical models of inflammation and demyelination and explore whether antagonized with the innovative aptamer ApTOLL would be beneficial throughout the course of pathology over time, as previously observed in other neurological diseases [16, 69].

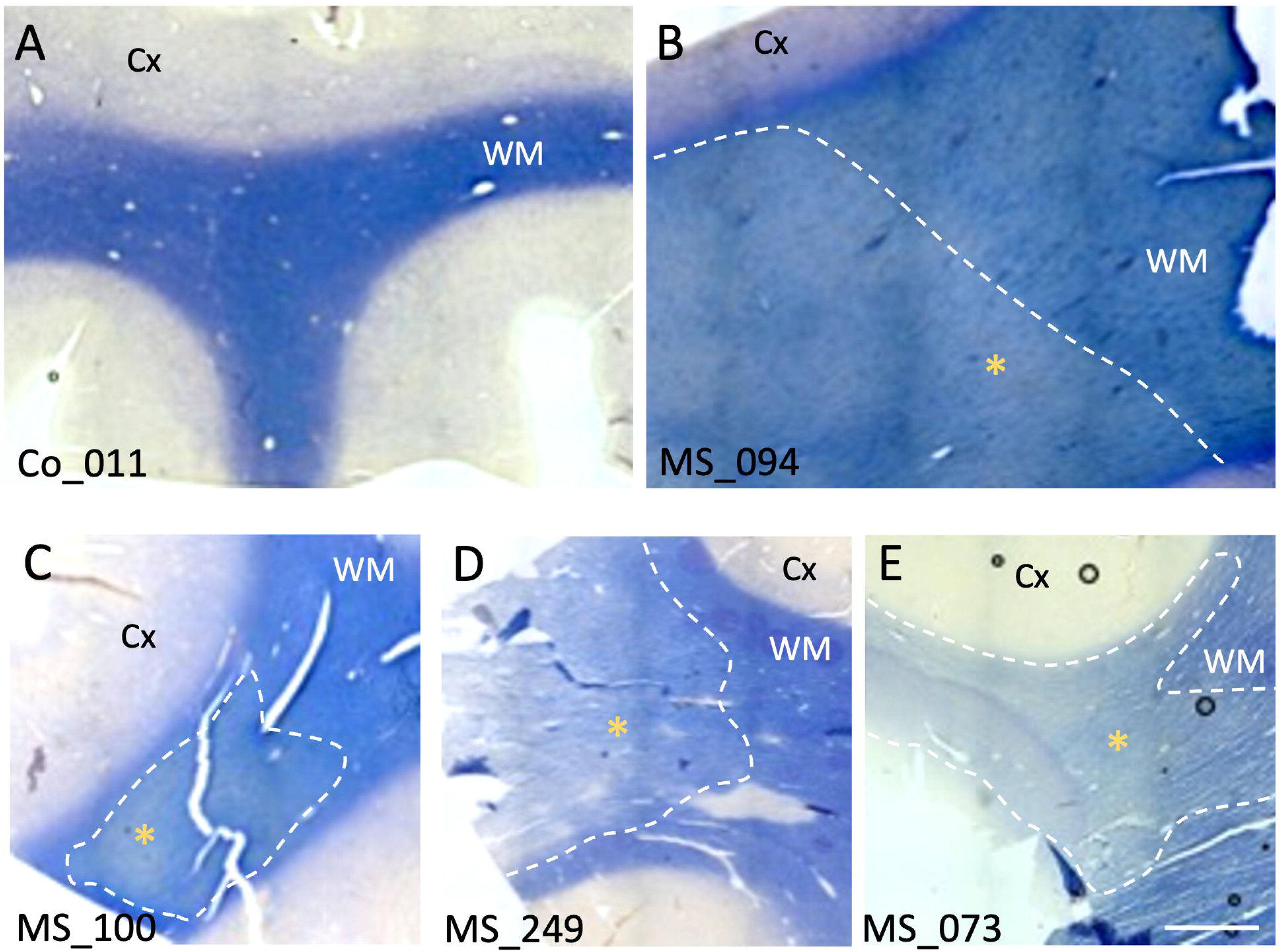

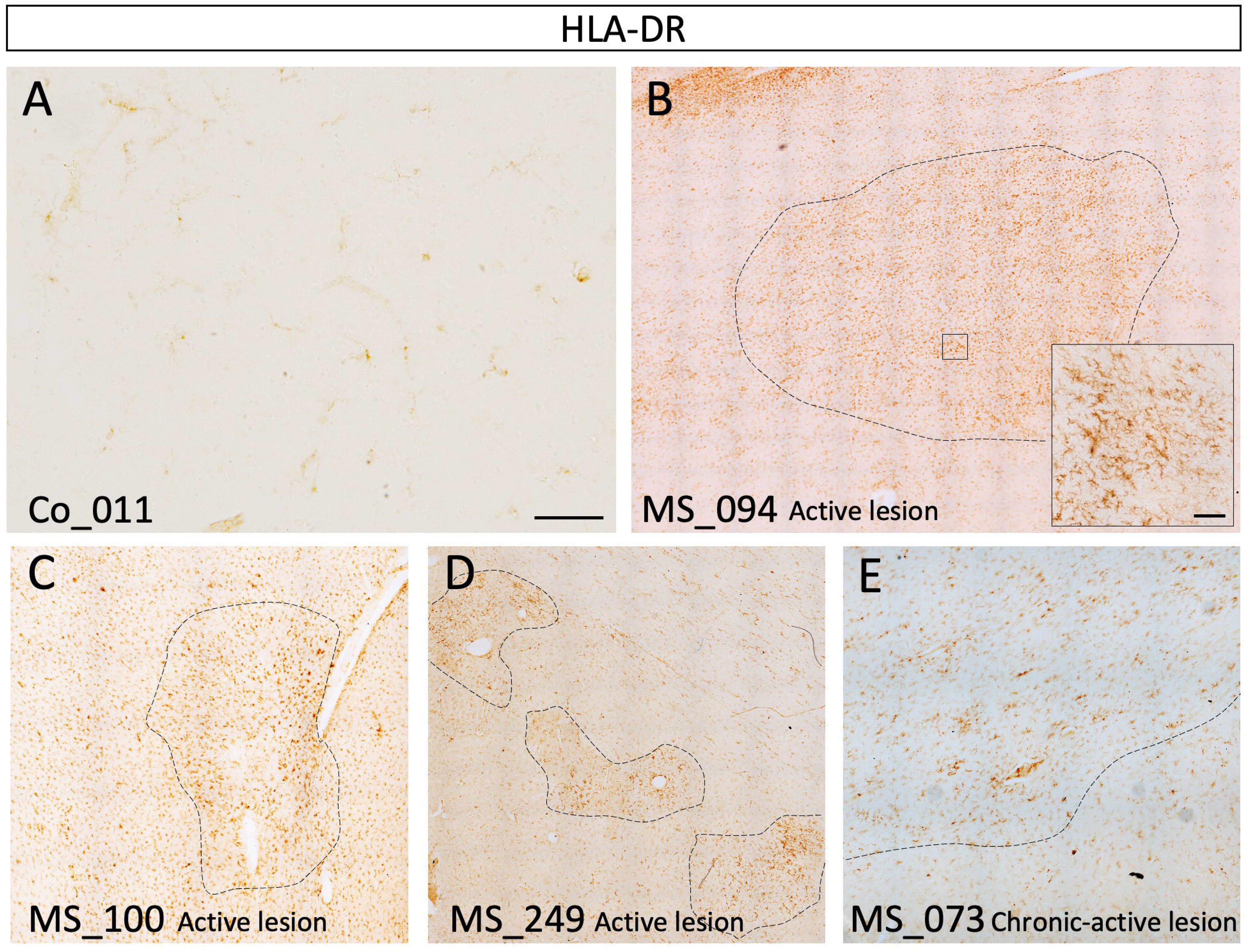

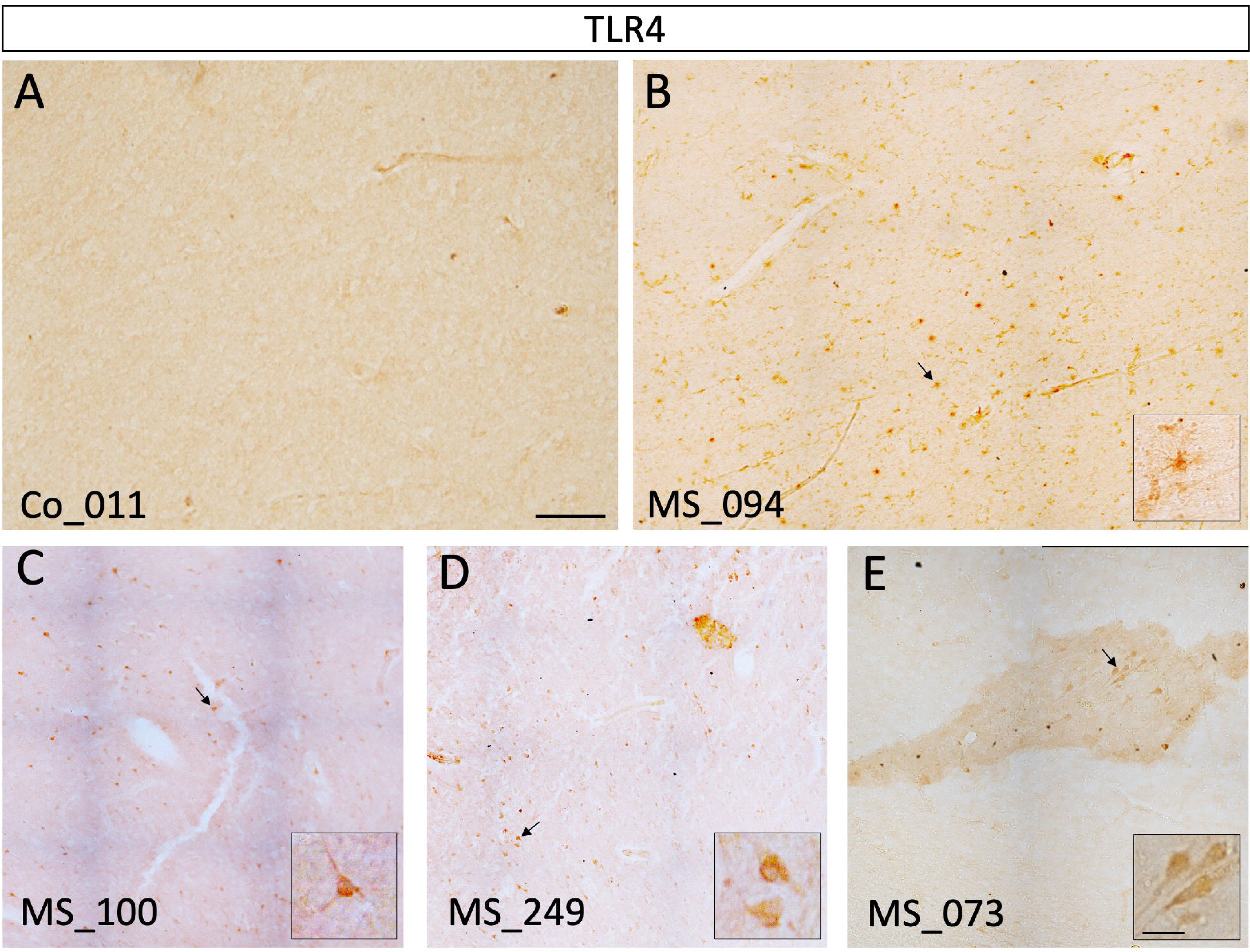

### ApTOLL treatment reduces EAE severity and demyelination in the spinal cord

In this model of EAE, there is T-cell driven inflammation and demyelination in the spinal cord, modeling the highly inflammatory early phase of MS. The optimal dose of ApTOLL for reduction of clinical score was determined by assessing four different concentrations (0.45 mg/Kg, 0.91 mg/Kg, 1.82 mg/Kg and 3.6 mg/Kg) administered through a single i.v. injection at the onset of EAE symptoms (see Methods section for details). The recovery of the clinical scores reached were significantly lower when the two intermediate doses were used (Supplementary Fig. 1A, online resource). Importantly, the control group did not present any pathological manifestations in these studies (data not shown). Histology study showed significantly less demyelination after administration of both the 0.91 and 1.82 mg/kg doses (Supplementary Fig. 1B-C, online resource), although the significantly better preservation of myelin (MBP staining) and axons (NFH staining), as well as the significantly larger proportion of OPCs (PDGFRα immunostaining) with no apparent changes in microglia (Supplementary Fig. 1D-G, online resource), led us to choose the dose of 0.91 mg/kg ApTOLL as optimal for our purposes (as used in the rest of our study). Similarly, we determined the therapeutic window for the single dose of ApTOLL, at the onset of EAE symptoms, 24h post-onset or at the peak of the symptoms. Although there was a reduction in clinical severity in each of these three treatment groups, the effects of ApTOLL were strongest when it was administered at the onset and 24h post-onset (Supplementary Fig. 2A, online resource). Demyelination was significantly lower in the three groups treated with ApTOLL when compared with the EAE-Veh animals (that received the vehicle alone), although this effect was particularly stronger when this agent was administered at the onset of EAE symptoms (Supplementary Fig. 2B- C, online resource), as it was in general for the rest of the histological parameters studied (Supplementary Fig. 2D-G, online resource). Together, our results suggest that a single dose of 0.91 mg/kg ApTOLL administered as soon as the mice presented symptoms was effective to protect the CNS from the demyelinating insult and that there was at least some evidence that spontaneous remyelination may also have been promoted.

With this selected dose, differences between the EAE-Veh and EAE-ApTOLL-treated group were observed from the beginning of the treatment, as there was a clear reduction in the slope of the curve and the disease scores obtained for the latter were significantly lower from day 4 onwards (Fig. 2A). The EAE-ApTOLL treated mice also showed a high recovery rate at the end-point compared to both the vehicle mice (67.2 ± 2.4%) and their own clinical scores at the peak of the disease (ApTOLL recovery at the end-point vs. peak: 61.8 ± 4.9%). In accordance with these results, EAE-ApTOLL-treated animals had fewer and relatively smaller demyelinated lesions in the WM (mainly ventro-lateral) than the EAE-Veh mice (Fig. 2B). Indeed, the total demyelination diminished significantly (by 75%) in the animals that received ApTOLL (Fig. 2C). The areas of well-preserved myelin and neurofilaments (expressing MBP and NFH, respectively) were significantly larger in the EAE-ApTOLL-treated animals than in those EAE-Veh animals (Fig. 2D-F). Although a significant increase in the number of cells in the oligodendroglial lineage (Olig2^+^ cells) was observed following ApTOLL treatment (Fig. 2G), the effects were particularly notable for OPCs (Olig2^+^/PDGFRα^+^ cells) and mature oligodendrocytes (Olig2^+^/CC1^+^ cells: Fig. 2H).

The aptamer specificity was also studied by different assays. On the one hand, the non-agonist effect of ApTOLL on all TLRs was characterized (Supplementary Fig. 3A, online resource). Furthermore, another experiment was carried out to enable early identification of significant off-target interactions with ApTOLL. Results showing an inhibition higher than 50% were considered to represent significant effects of the test compounds. Such effects were not observed at any of the targets studied (Supplementary Fig. 3B, online resource). Despite the values obtained in the enzyme assays did not show significant effects, a deep characterization was performed for PDE3A and PDE4D2 and no significant results were obtained (Supplementary Fig. 3C, online resource). Therefore, no inhibitory effect was detected in any target selected.

### In murine EAE, TLR4 is mostly expressed by Iba1^+^ cells and its blockade decrease inflammation

As inflammation is one of the main characteristics of MS and due to the central role of TLR4 in the activation of microglia, we determined the presence of Iba1 ^+^ cells cells in the spinal cord of EAE mice that had been injected with ApTOLL at the onset of symptoms. A significant decrease to approximately half the number of these inflammatory cells was detected, especially in the lesion areas (Fig. 5A-B). Similarly, the quantitative analysis of demyelinating lesions showed a significant reduction in the density of TLR4-labeled cells after treatment with ApTOLL (Fig. 5A, C), as well as in those cells co-expressing both markers (Iba1 ^+^/TLR4^+^ cells: Fig. 5A, D). The number of both TLR4+ and Iba1+ cells significantly correlated (Fig. 5E). To identify different microglia/macrophages states within demyelinating lesions of EAE mice, we used CD68 to recognize microglia/macrophages reactive to damage, and CD206 to identify a maintenance/repair phenotype [62]. We observed that ApTOLL-treated animals expressed significantly less microglia/ macrophages in a reactive state but more in a reparative state (Fig. 5F-H). Moreover, this process that seemed to be favored by the shift of these cells towards repair phenotypes, was evident through their morphology (larger number and longer length of branches) [9, 27, 61], after exposure to a demyelinating insult and the subsequent treatment with ApTOLL (Supplementary Fig. 4, online resource).

### ApTOLL also promotes neuroprotection and myelin preservation in the murine cuprizone model of MS

To better evaluate the effects of ApTOLL on oligodendroglial biology, we took advantage of the CPZ *in vivo* model as this copper chelator induces reversible demyelination in the CNS, particularly in the CC [38]. Mice were maintained on a CPZ diet for 6 weeks with a weekly dose of ApTOLL throughout (see Materials and Methods for details; Fig. 6A). While inflammation (Supplemmentary Fig. 5A-B) and astrogliosis (Supplemmentary Fig. 5C-D) significantly decreased in the ApTOLL-treated animals, OPC proliferation at the end of week 3 (by quantifying the co-immunolabeling of Olig2 -oligodendroglial lineage- and EdU incorporation -proliferating cells-; [25]) was significantly promoted (Supplementary Fig. 5E-F, online resource). After 6 weeks with cuprizone diet, when the peak of demyelination occurs [32], animals were tested in the rotarod: the motor coordination as well as the latency to fall were severely affected in the CPZ-Veh mice when compared to the control mice, but the administration of ApTOLL improved their motor function, these closely resembling the control animals (Fig. 6B). In agreement with previous reports [26, 49, 75], a six-week diet with cuprizone induced almost complete loss of myelin in the CC in the CPZ-Veh group, while weekly ApTOLL injection partially but significantly protected this tissue against demyelination (Fig. 6C-D). The role of this compound in the preservation of mature myelinated axons was also suggested by the number of nodes of Ranvier: while this was markedly reduced in CPZ-Veh demyelinated animals, ApTOLL treatment doubled their number, although remaining significantly far from the control group (Fig. 6C, E). In addition, the number of OPCs significantly recovered after ApTOLL administration, and although the number of mature oligodendrocytes significantly doubled with the treatment, they remained lower (just about one fourth) than in the control mice (Fig. 6C, F-G).

### ApTOLL treatment enhances OPC proliferation and differentiation *in vitro* without affecting their survival

The *in vivo* data obtained previously in these animal models of MS suggest that ApTOLL affects different pathogenic aspects, maybe including the promotion of spontaneous remyelination (Figs. 4, 5 and 6). To further elucidate whether ApTOLL would exert a direct effect on OPCs, we verified the expression of TLR4 in these cells. This receptor was expressed by aprox. 40% of isolated OPCs (Fig. 7A; Supplementary Fig. 6A-B), and it was not affected by inflammatory conditions (IL1-β or TNF-α; Supplementary Fig. 6A-B, online resource). In addition, ApTOLL binding to TLR4 was proved by tagging the aptamer with an Alexa-488 molecule (Fig. 7B). These data strongly suggest that ApTOLL would also directly modulate OPC biology.

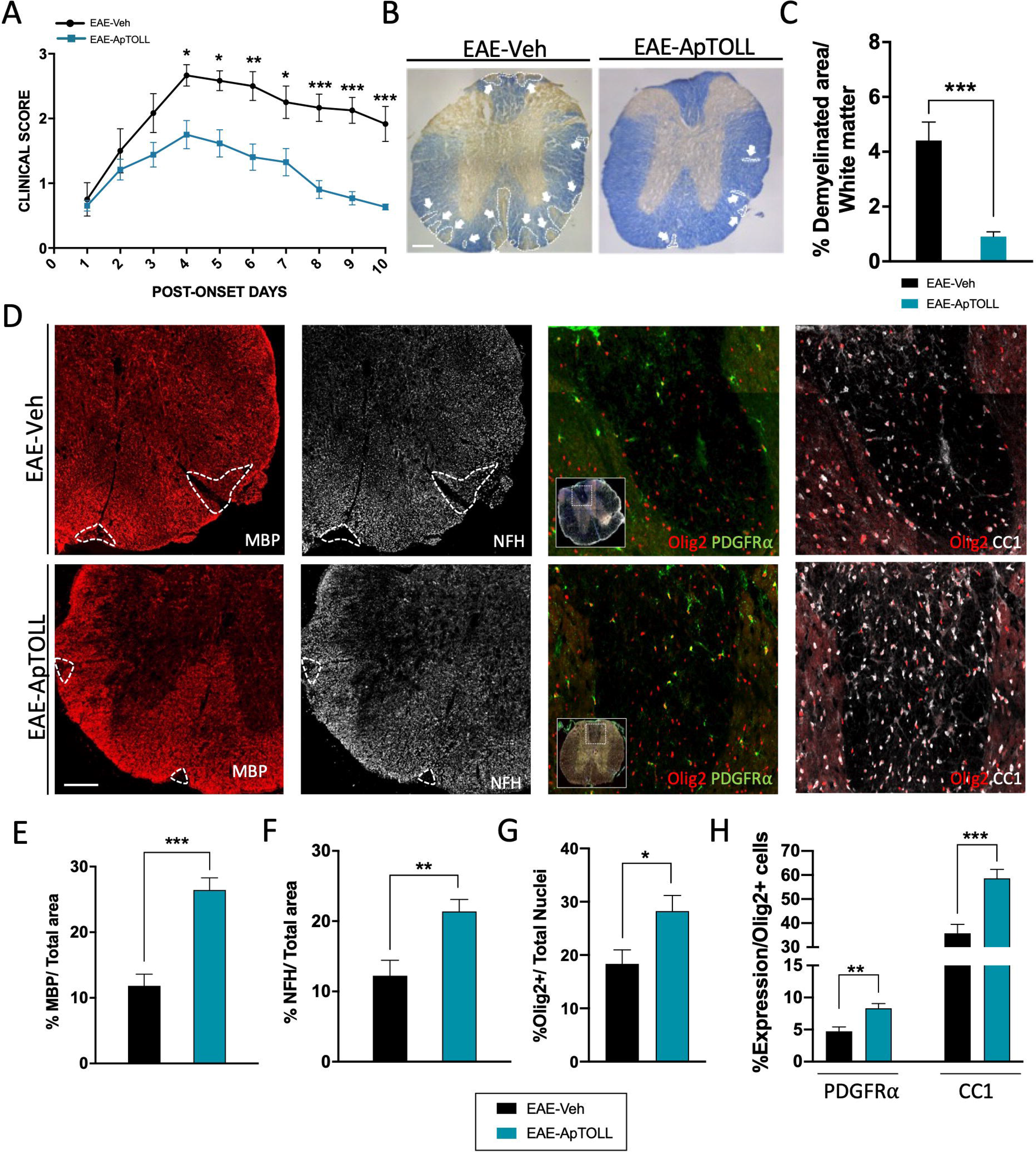

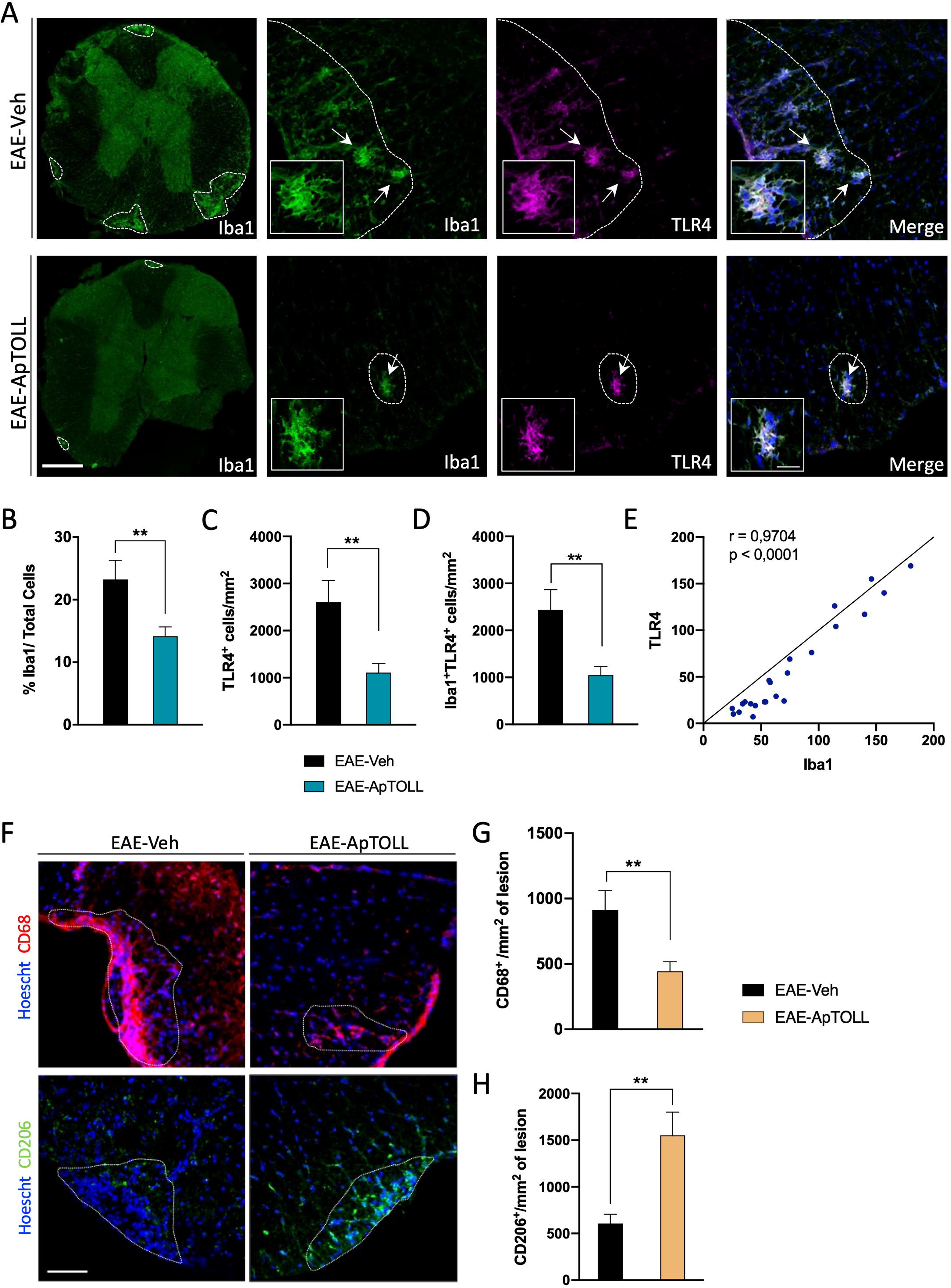

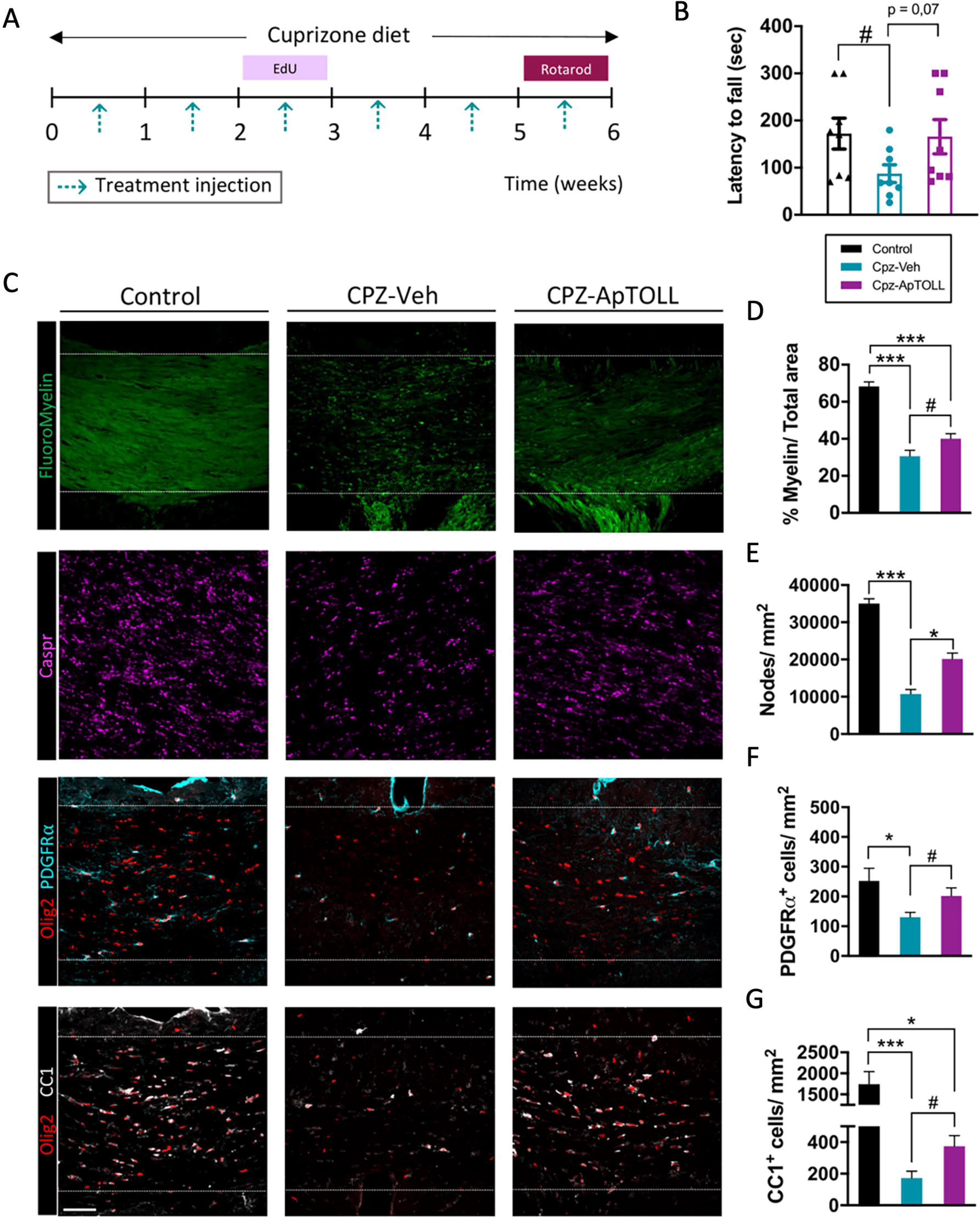

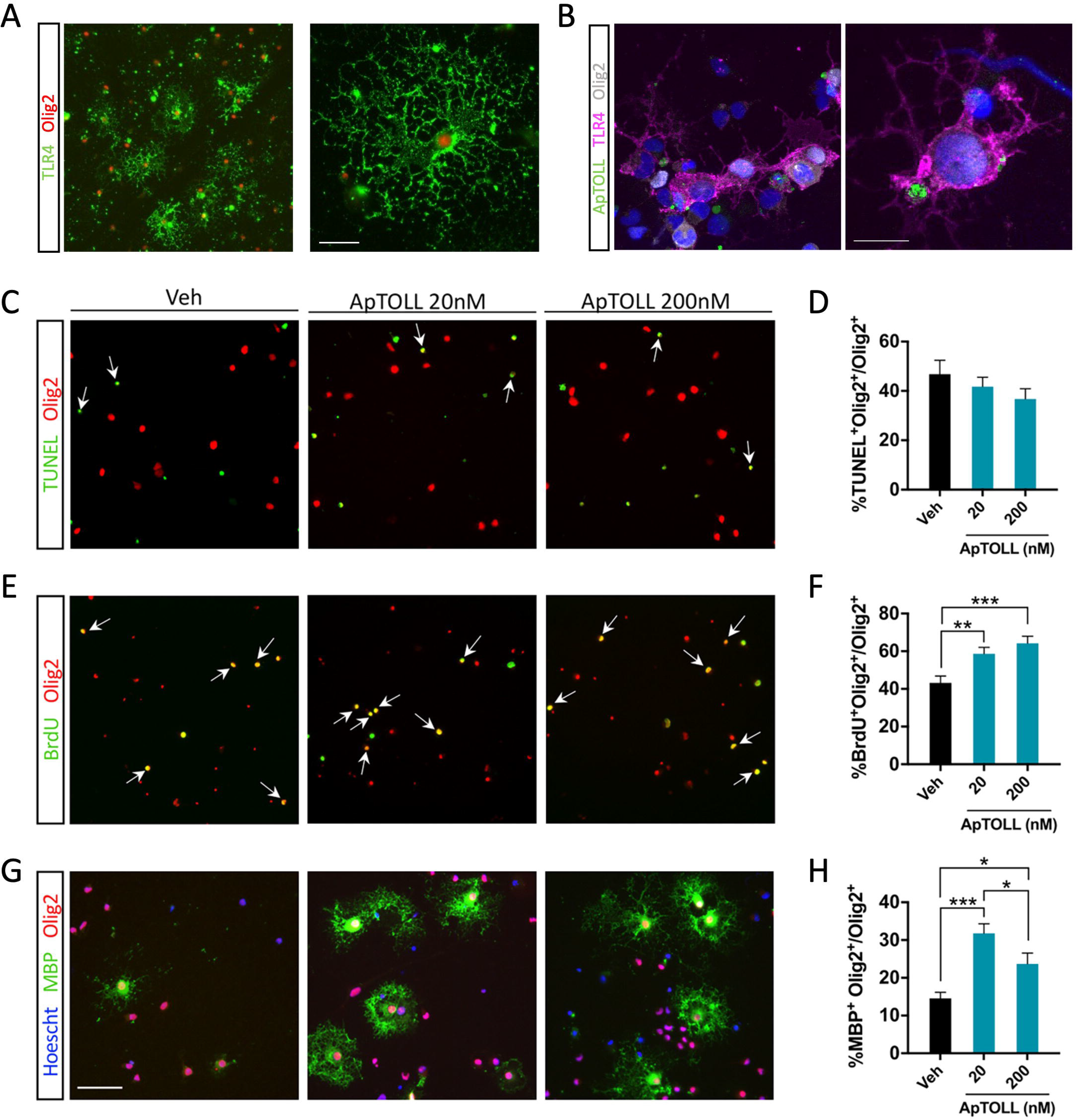

We also tested the effects of the aptamer at two different concentrations (20 nM and 200 nM, as tested previously: [16]) in the survival, proliferation and differentiation of OPCs in purified primary cultures. Neither OPC viability in MTT assays (data not shown) nor their survival in TUNEL assays (Fig. 7C-D) were affected by ApTOLL, however exposure to this aptamer did significantly induce OPC proliferation (Fig. 7E-F). Regarding the maturation of OPCs towards myelinating phenotypes and their myelin production, we maintained cells in culture for 5 days under the different experimental conditions. Exposure to ApTOLL produced a significant increase not only in the number of mature oligodendrocytes expressing the MBP marker (Fig. 7G-H) but also, in the area occupied by this antibody relative to the control OPCs (MBP^+^ area in Veh 4711 ± 1402 μm^2^, in ApTOLL [20 nM] 8082 ± 757 μm ^2^ and in ApTOLL [200 nM] 9871 ± 1415 μm^2^: One-Way ANOVA Dunn’s *post hoc* test Veh vs. ApTOLL [20 and 200 nM] p < 0.001). Similar results were obtained in murine OPCs (data not shown). As an additional proof of TLR4 mediation in these effects, we replicated these experimental series in OPCs isolated from the brain cortex of neonatal TLR4^-/-^ mice, and the exposure to ApTOLL or the vehicle alone did not induce modifications in any OPC survival, proliferation or differentiation (Supplementary Fig. 6C-H, online resource). Accordingly, we concluded that ApTOLL exerts a direct promyelinating role *in vitro* and that this seems to be driven through TLR4 expressed by oligodendroglia.

### ApTOLL promotes remyelination *ex vivo*

To study the impact of ApTOLL on myelin regeneration after a demyelinating insult, *ex vivo* organotypic cerebellar slice cultures were prepared from neonatal mice and tissue damage was provoked with LPC, a membrane-disrupting chemical that provokes rapid loss of myelin and oligodendrocytes [3, 54] (Fig. 8A). Demyelination with LPC was confirmed after being removed from the medium with a 55-60% reduction in both the amount of myelin and its co-localization with neurofilaments. To study the remyelinating effect of the aptamer, we maintained the slices for 6 dpl and increased myelin production was evident in LPC slices that received ApTOLL than in those that received the vehicle alone (Fig. 8B-C). Moreover, treatment with ApTOLL resulted in a significantly higher proportion of remyelinated axons than when they were exposed to the vehicle alone (Fig. 8B, D), the latter only showing a 22% increase over the control at 0 dpl (LPC+Control 22.6% ± 3.0, LPC+Veh 27.6% ± 6.8: Student’s *t*-test p = 0.45), whereas exposure to ApTOLL produced a 130% increase (LPC+Control 22.6% ± 3.0, LPC+ApTOLL 51.8% ± 6.7: Student’s *t*-test p < 0.001). These data confirmed the ability of ApTOLL to enhance new myelin generation and promote the formation of new sheaths around axons that had been previously demyelinated.

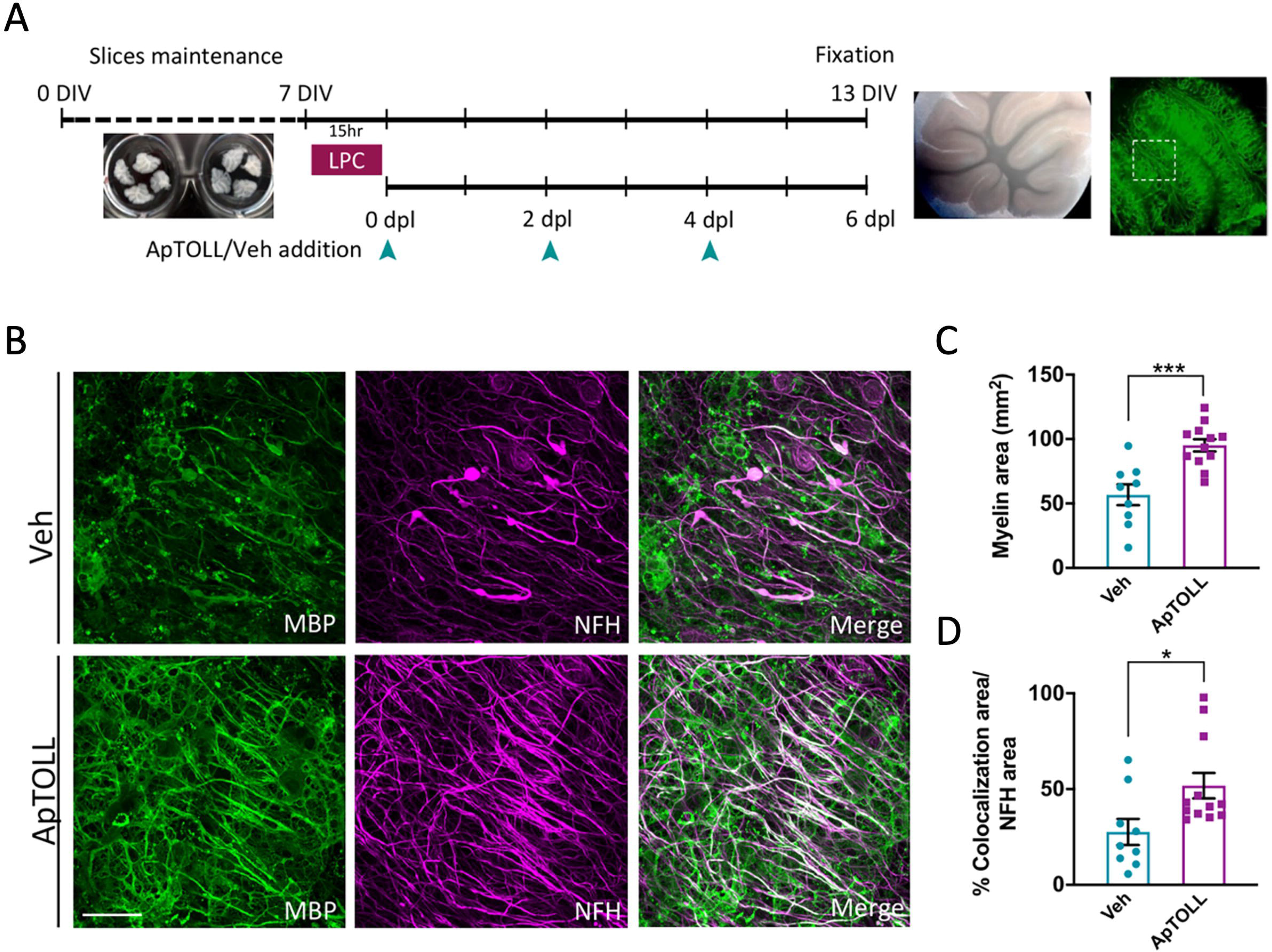

### ApTOLL enhances the maturation of human adult OPCs towards myelinating phenotypes and increases myelin production

Based on the data obtained from rat OPCs *in vitro*, we determined whether ApTOLL exerted a promyelinating effect in primary cultures of adult human OPCs isolated from non-tumor biopsies of the adult cerebral cortex (see Materials and Methods). As described previously [6, 50, 51], it was necessary to maintain these cells for at least 15 DIV to differentiate them. Exposure to ApTOLL (20 nM) gave rise to significantly more mature oligodendrocytes (CC1 ^+^) than when the OPCs were cultured in control conditions, and these differences were even more pronounced when 200 nM ApTOLL was used (Fig. 9A, B). However, both these treatments gave similar results in terms of the myelin proteins generated by these mature oligodendrocytes (measured as MBP ^+^ area) with respect to the total number of cells quantified in each field (Fig. 9A, C). Together, these results show that as seen with rodent cells, ApTOLL may have a direct effect on promoting the production of proteins needed for myelin formation by human adult OPCs and they suggest that ApTOLL may be a suitable agent to directly promote effective remyelination in MS in addition to its anti-inflammatory activities.

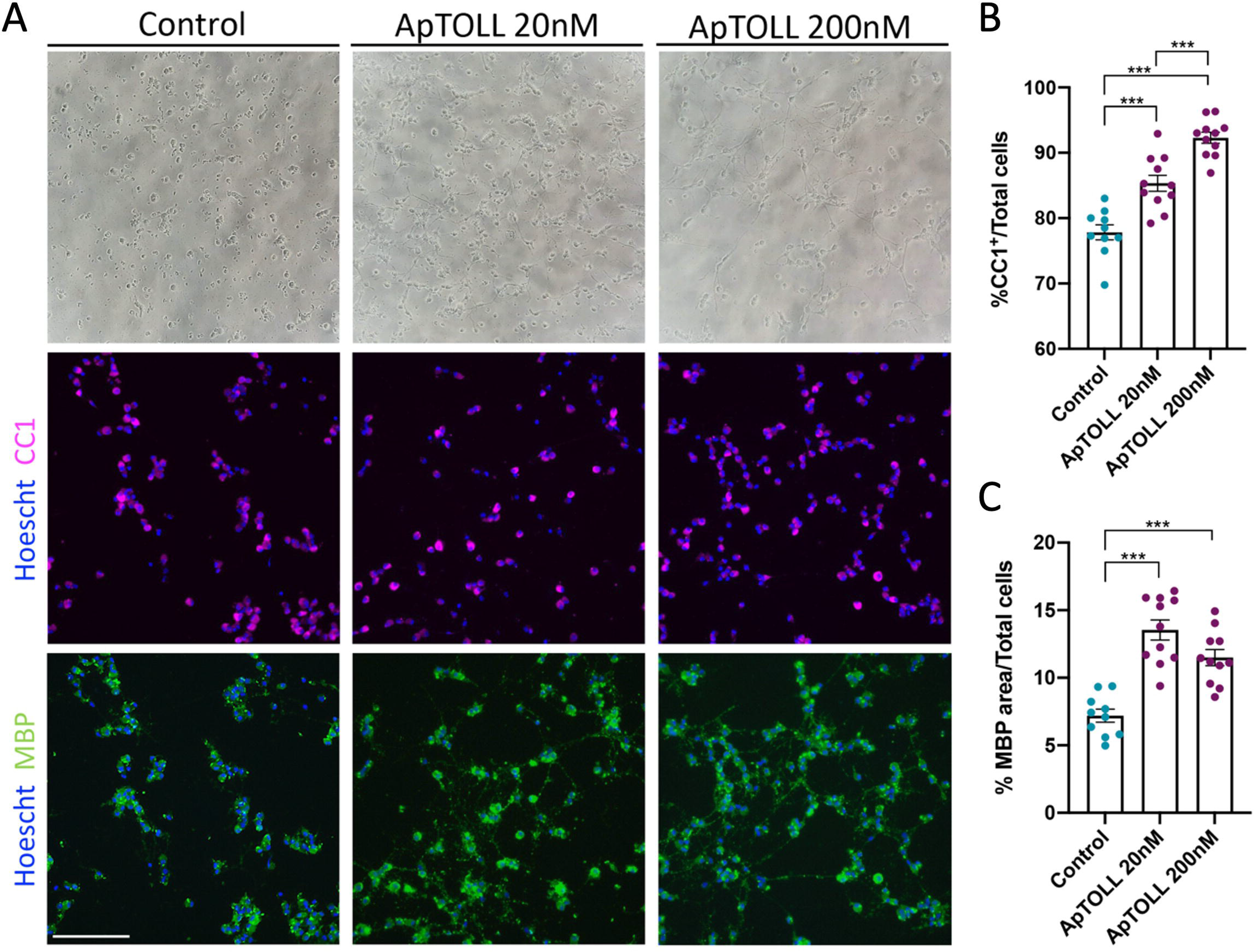

## DISCUSSION

The contribution of the inflammatory-immunological component to MS pathogenesis has been extensively demonstrated, and it is known that TLRs (including TLR4) play important roles [1, 22, 60, 83]. Nevertheless, and up to our knowledge, our present work is the first to identify TLR4 in human MS demyelinating lesions. The new selective TLR4 antagonist aptamer (ApTOLL) is subject of clinical trials to treat cerebral stroke and other diseases [30, 31, 63, 69], and here we propose that it could also be a promising candidate for the treatment of MS. Indeed, we not only reveal a possible neuroprotective activity of this antagonist by limiting the inflammatory response but also demonstrate its direct incorporation to OPCs and ApTOLL promotion of OPC differentiation and myelin formation.

Immunohistochemistry showed TLR4-expressing cells in demyelinating human MS lesions. This result had not yet surfaced despite previous studies such as those based on the RNAseq technology [15, 78]. The distribution of TLR4-positive elements varied in the different histopathological type of lesions: while they were abundant in active lesions (even more than the abundant inflammatory infiltrating cells), these elements drastically reduced their number in the lesions with chronic aspect. Furthermore, the morphology of TLR4-positive elements also varied according to the type of lesion: in active lesions, cells appear to have a more rounded soma or a pyramidal shape, and generally few branches; in chronic lesions, this morphology suggested cells of a neural nature, very similar to the FGFR1 ^+^-OPCs in the periplaque of chronic-active lesions [8]. However, in our present work the TLR4 ^+^-cells were almost circumscribed to the core of chronic-active demyelinating lesions.

Based on the presence of TLR4-positive cells in these lesions of MS patients, we evaluated the potential to use an aptamer to inhibit the activation of the TLR4 signaling. Aptamers are considered candidates for inclusion in therapeutics due to their high affinity and specificity, as well as their advantages over conventional drugs or antibodies (safety, stability, *in vitro* selection, ability to be modified without structural alterations, production costs, etc) [35, 36]. In the present study we have used ApTOLL, an aptamer specifically developed to antagonize TLR4. This aptamer has been shown to be effective in other models of pathology such as ischemic stroke and myocardial infarction [16, 63, 69].

Here we also show how an ApTOLL single injection resulted in a significant recovery of clinical symptoms in the EAE model, which was associated with increased myelin and axonal preservation. The aptamer notably reduced the number of microglial and TLR4-expressing cells, which could be due to i) the blockage of the membrane binding site and thereby preventing its activation by other ligands, or to ii) the internalization of the receptor once it is antagonized, as proposed previously [64, 81]. The correlation between the activation of this receptor and microglia/macrophages behavior might suggest that TLR4 blockade dampens microglial activation and, therefore, the inflammatory process [42, 85]. However, those cells that remain active could be participating in phagocytic functions of myelin or axonal debris [68], which is also considered as a necessary process for proper remyelination [18, 46]. Hence, the specific antagonism of TLR4 by ApTOLL but not its complete depletion may produce a remarkable clinical improvement, related to the modulation of an excessively strong immune response, preventing tissue damage. Interestingly, the therapeutic use of TLR4 humans is currently being studied in clinical trials in acute ischemic stroke (NCT04734548) and COVID-19 (NCT05293236) patients and has recently demonstrated good safety and pharmacokinetic properties in healthy volunteers [31].

The role of ApTOLL does not seem to be reduced to its effect on microglia cells and thus on the inflammatory component. In this sense, we have observed a remarkable effect on the elements involved in the remyelination process, i.e. on OPCs. This effect goes beyond the fact that ApTOLL decreases inflammatory conditions, to which elements of the oligodendroglial lineage are particularly sensitive [41, 42, 73, 85]. In a non-inflammatory demyelinating scenario, the cuprizone model, the treatment with ApTOLL significantly promoted the number of OPCs (better survival plus promoted proliferation) and mature oligodendrocytes and myelin distribution. All together could be reflecting acceleration of spontaneous remyelination. This was corroborated in OPC primary cultures, where we confirmed the expression of TLR4 and the direct binding of ApTOLL to these cells. We can conclude that this pro-myelinating effect of the aptamer would not be minor since it is known that for successful remyelination quiescent OPCs must actively proliferate and properly recruited towards demyelinating areas [5, 13, 57] (for recent reviews on this specific subject, see [20, 71]).

Nevertheless, it is not only important to increase the number of OPCs, but clinical benefits rely on their differentiation towards myelin-forming phenotypes. It has been argued that the proliferation and differentiation of OPCs are sequential events that cannot take place simultaneously [65], however these results are in line with others that also show that the proliferation of OPCs and the maturation process can coexist [28, 54, 55, 79, 82]. The present data from human adult OPCs strongly support that this would also be the case on human MS. In spite of the remarkable heterogeneity of OPCs recently demonstrated [6, 33, 34, 48], the present observations suggest that ApTOLL effects on these cells do not vary according to age or species, which would be very relevant for the future design of clinical trials.

We also showed that the treatment with ApTOLL promoted the coating of axons that had been demyelinated in organotypic cultures, then favoring remyelination. Previous data from various authors indicate that the Wnt/β-Catenin and Akt/mTOR pathways play an important role during the differentiation of OPCs and the myelination process [11, 58]. It has been shown that there is an interaction and mutual regulation between these signaling pathways and NF-κB, a product of the TLR4 pathway [12, 21, 47]. One possible explanation is that, in oligodendroglia, not being inflammatory cells, TLR4 activation/inhibition could be affecting other signaling pathways with which molecules involved in the TLR4 pathway interact, such as Wnt/β-Catenin, Akt/mTOR or ERK/MAPK. However, this field is still unknown, and it would be very interesting to study the molecular mechanisms by which TLR4 can act in non-inflammatory cells such as oligodendrocytes. Together, our current findings suggest a new therapeutic approach for the treatment of inflammatory and demyelinating diseases like MS. The molecular nature of the aptamer, ApTOLL, specifically blocking TLR4 exerting a clear anti-inflammatory effect as well as an indirect/direct neuroprotective and remyelinating effect confers advantages over other compounds, and also has a good safety profile, as demonstrated in the First-in-Human study [31]. This opens the door to future clinical studies with this aptamer in MS human trials.

## Acknowledgements

This work was supported by grant IND2018/BMD-9751 (Programa de Doctorados Industriales, Comunidad de Madrid, Spain), SAF2016 -77575-R (Spanish Ministerio de Economía, Industria y Competitividad-MINECO), and the contract for technological support ApTLR2019-PC-MS-001 (AptaTargets, S.L., Spain) to FdC. BF-G is currently hired by Aptatargets S.L., PG-M is hired under PEJ-2020-AI/BMD-18541 de la Comunidad de Madrid, Spain (associated with the youth guarantee fund to FdC), SN had a predoctoral contract from the UCLM and was hired under SAF2012 -40023, SAF2016- 77575 -R, RD12 -0032/0012 and RD16 -0015/0019 (Spanish Ministerio de Economía, Industria y Competitividad-MINECO) and IND2018/BMD-9751, YL has been contracted under ReTics and SAF (to FdC). We thank David Segarra and Ma Eugenia Zarabozo (AptaTargets S.L.) for their constant technological support, Laude Garmendia for her indispensable constant help at the animal facility (Instituto Cajal-CSIC), including the extra effort during Covid-19 pandemics, Profs María Ángeles Moro (Centro Nacional de Investigaciones Cardiovasculares Carlos III, Madrid and Universidad Complutense de Madrid, Spain) and Ignacio Lizasoaín (Universidad Complutense de Madrid, Spain) for lending us the TLR4 knockout mice, and the former GNDe member Dr. Carolina Melero- Jerez (currently working at JazzPharma, Spain) for the initial training of BF-G on EAE animal model and different techniques at the laboratory. Human samples were supplied by the UK Multiple Sclerosis Tissue Bank, funded by the Multiple Sclerosis Society of Great Britain and Northern Ireland (registered charity 207495).

