## Supplementary files for "ApTOLL, a new therapeutic aptamer for cytoprotection and (re)myelination after Multiple Sclerosis"

**Tables**

**Table 1.** Summary of the human samples analyzed.

| Patient | Age (years)/ Sex | Diagnostic | | TP  (h) | Disease duration (years) | Lesion activity  Active Chronic | Cause of death |
| --- | --- | --- | --- | --- | --- | --- | --- |
| MS60 | 55/M | SP | 16 | | 43 | 3 1 | Aspiration of gastric contents, MS |
| MS73 | 80/F | SP | 20 | | 50 | 2 3 | Bronchopneumonia |
| MS94 | 42/F | PP | 11 | | 6 | 4 3 | Bronchopneumonia, MS |
| MS100 | 46/M | SP | 7 | | 8 | 1 5 | Pneumonia |
| MS115 | 76/F | SP | 21 | | 58 | 2 1 | MS |
| MS125 | 76/F | SP | 13 | | 30 | 1 3 | MS |
| MS249 | 69/F | RR | 8 | | 42 | 3 1 | Thoracic infection,  heart failure |
| MS274 | 56/M | RR | 8 | | 19 | 2 0 | Cancer of oesophagus |
| CO02 | 85/M | Normal | 12 | |  |  | Prostate cancer |
| CO05 | 95/F | Normal | 10 | |  |  | Bronchopneumonia |
| CO11 | 77/M | Normal | 26 | |  |  | Carcinoma of the lung metastasised |
| CO13 | 73/F | Normal | 59 | |  |  | Cancer of oesophagus |
| CO25 | 35/M | Normal | 22 | |  |  | Carcinoma of the tongue |

Abbreviations: MS, Multiple Sclerosis; Co, Control; F, female; M, male; SP, secondary progressive; PP, primary progressive; RR, relapsing remitting; TP, time post-*mortem.*

| **Use** | **Antibody** | **Target** | **Dilution** | **Species** | **Supplier** |
| --- | --- | --- | --- | --- | --- |
| IHC | NFH | Neurofilaments | 1:1000 | Rabbit | Abcam |
| IHC | Iba1 | Microglia | 1:500 | Guinea pig | Synaptic system |
| IHC | CC1 | Mature oligodendrocytes | 1:200 | Mouse | Millipore |
| IHC | PDGFRα | OPCs | 1:200 | Goat | R&D systems |
| IHC | Caspr | Paranodes | 1:900 | Rabbit | Abcam |
| IHC | TLR4 | Toll-like receptor 4 | 1:100 | Rat (*mice samples*)/  Mouse (*human samples*) | Santa Cruz/  Abcam |
| IHC | HLA-DR | Human leukocyte antigen | 1:900 | Mouse | ThermoFisher |
| IHC | GFAP | Astrocytes | 1:500 | Mouse | Millipore |
| IHC | CD68 | Macrophages and microglia with phagocytic activity | 1:250 | Mouse | ThermoFisher |
| IHC | CD206 | Anti-inflammatory microglia | 1:200 | Goat | ThermoFisher |
| IHC/ICC | MBP | Myelin | 1:500 | Rat | Serotec |
| IHC/ICC | Olig2 | Oligodendroglial lineage | 1:200 | Rabbit | Millipore |
| ICC | BrdU | Proliferating cells | 1:1000 | Rat | Abcam |

**Table 2.** List of primary antibodies used

Abbreviations: IHC, immunohistochemistry; ICC, immunocytochemistry; OPCs, oligodendrocyte precursor cells.

Figures

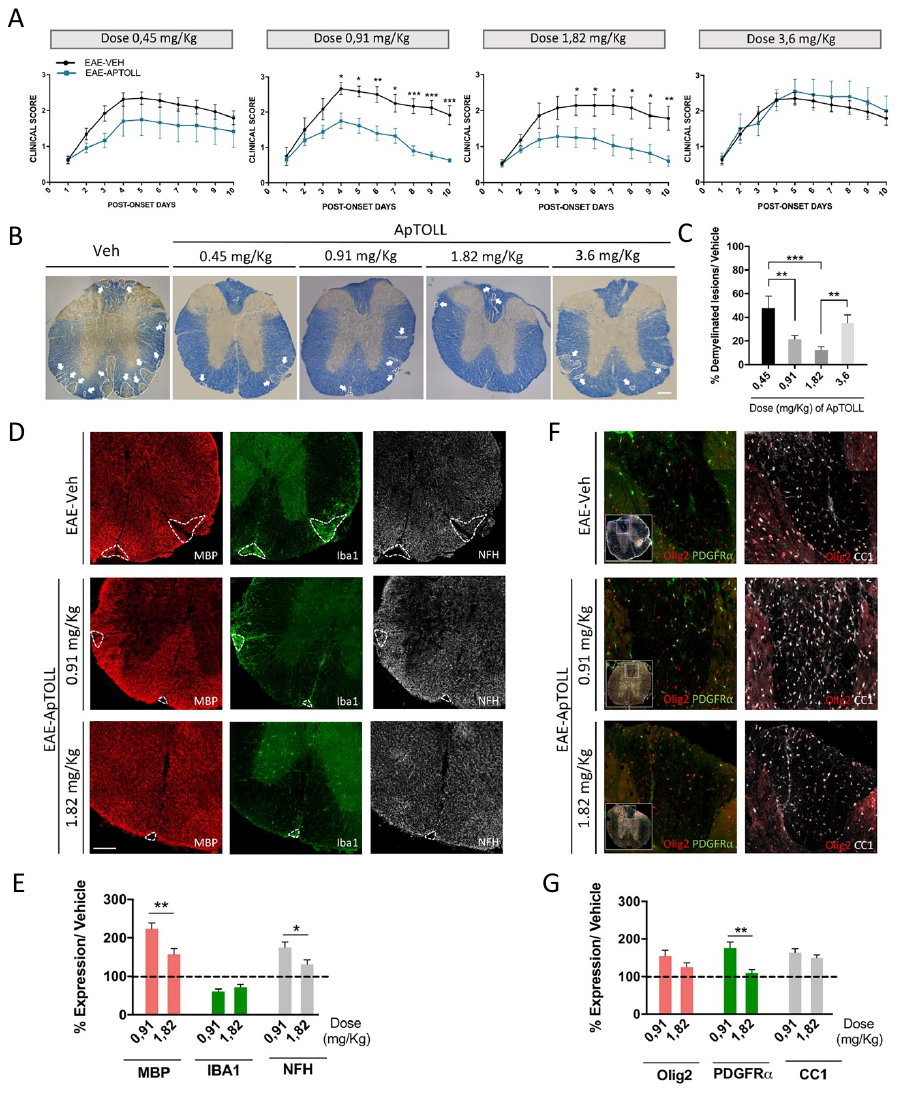

**Supplementary Fig 1. ApTOLL dose-response study in EAE. A:** Clinical course of the animals treated with each dose of ApTOLL compared to each vehicle group. There is a significant recovery in the clinical course of EAE after the injection of ApTOLL at the time of the onset of symptoms, both at a dose of 0.91 mg/Kg and 1.82 mg/Kg. **B-C:** Demyelinated areas are indicated with white arrows (**B**) and quantified with respect to the white matter area in each experimental group (**C**). The optimal doses for ApTOLL treatment in the murine EAE model of MS are the intermediate ones. **D-E:** Images of the different markers in the lesions (dashed white lines): MBP (red), NFH (gray), and Iba1 (green) (**D**). Graphs representing the recovery rate of ApTOLL-treated mice relative to the vehicle group (**E**). **F-G:** Magnified images (**F**) and histograms (**G**) of the oligodendrocyte studies at different stages of maturation. The 0.91 mg/Kg dose is optimal for ApTOLL-treatment in EAE. Scale bar, 200 μm in B, D and F. EAE-ApTOLL n=6, EAE-Veh n=4, Control n=5 for a dose of 0.45 mg/kg, EAE-ApTOLL n=13, EAE-Veh n=6, Control n=10 for a dose of 0.91 mg/kg; EAE-ApTOLL n=8, EAE-Veh n=7, Control n=8 for a dose of 1.82 mg/Kg; EAE-ApTOLL n=5, EAE-Veh n=4, Control n=5 for a dose of 3.6 mg/Kg. Results of the one-way ANOVA with a Tukey’s *post hoc* test and of the Student’s *t*-test for two independent groups are represented as: *p <0.05, **p <0.01, and ***p <0.001.

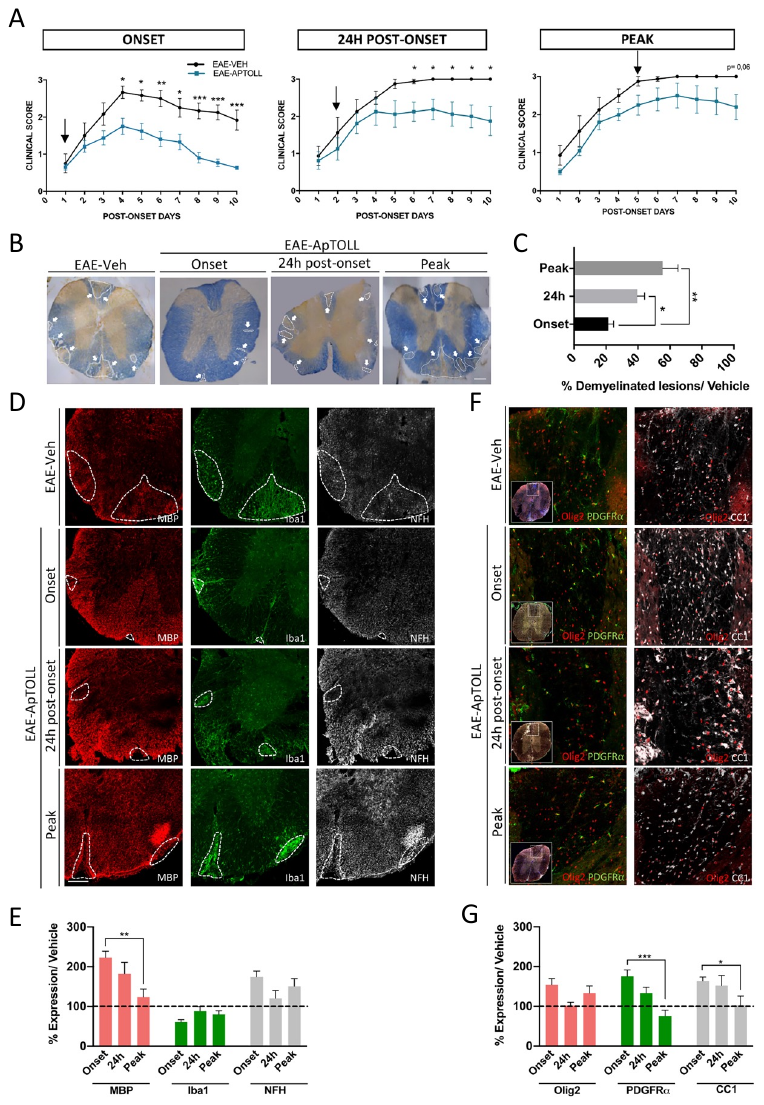

**Supplementary Fig 2. Determination of the therapeutic window with the optimal dose of ApTOLL**. **A:** Clinical course of the animals treated with ApTOLL at different time points of the disease relative to their respective vehicle group: onset (EAE-Veh n=6; EAE-ApTOLL n=13), 24h post-onset (EAE-Veh n=4, EAE-ApTOLL n=4) and peak (EAE-Veh n=4, EAE-ApTOLL n=5). **B-C:** Representative images of EC staining (**B**) and the quantification of demyelination relative to their respective vehicle group (**C**). White arrows indicate lesions. The treatment is more effective when injected at the onset of symptoms. **D-E:** Representative images (**D**) and graphical representation of the recovery rate of MBP, NFH and Iba1 relative to the vehicle group (**E**). **F-G:** Representative magnification of the oligodendrocyte labeling at different stages of maturation (**F**) and a graph of the oligodendrocyte ratio at different times of injection (**G**). Scale bar, 200 μm in B, D and F, and the results of the one-way ANOVA with a Dunn’s or Tukey’s *post hoc* test are represented as: *p <0.05, **p <0.01, and ***p <0.001.

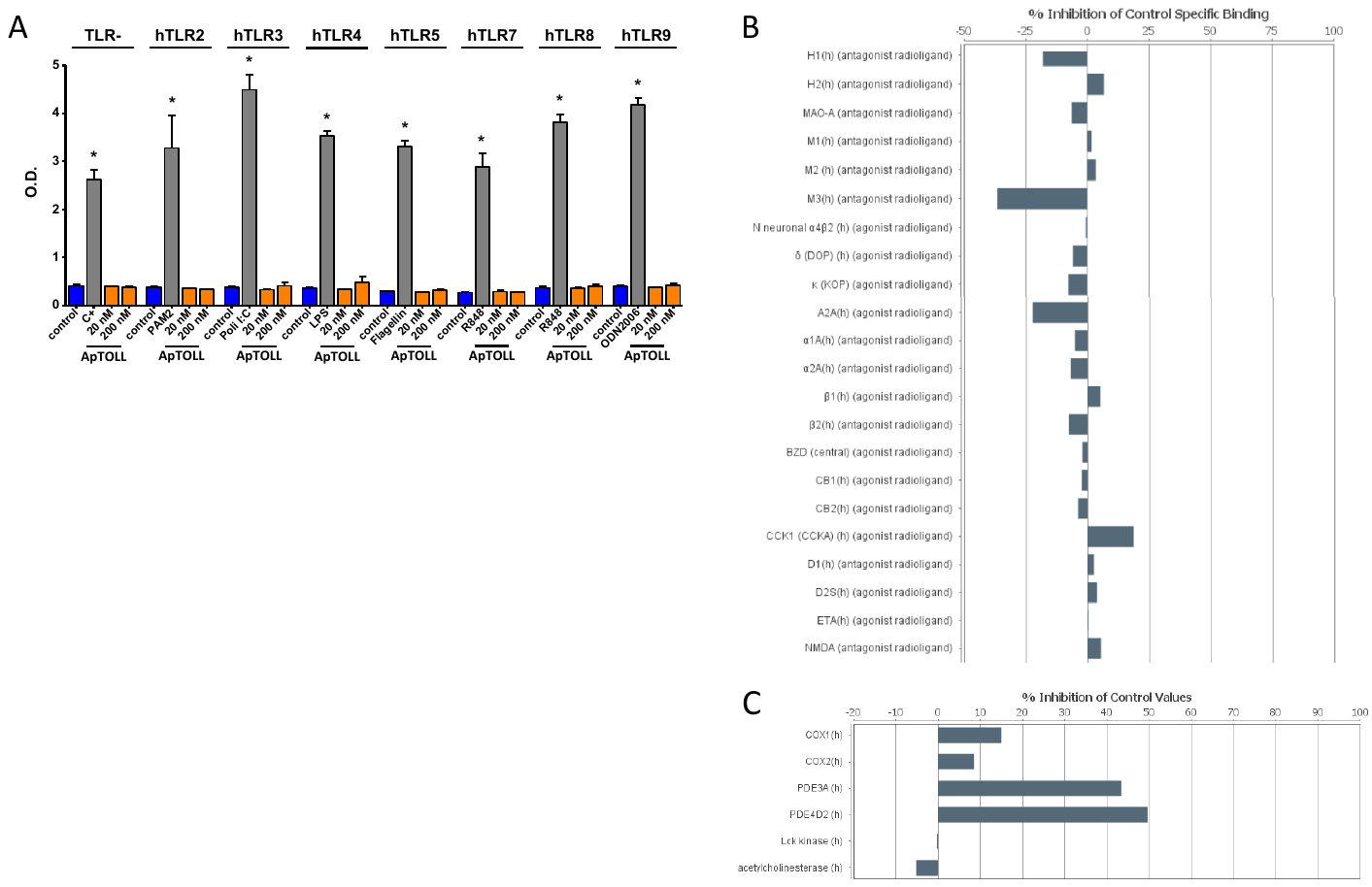

**Supplementary Fig 3. Study of ApTOLL specificity. A:** No agonistic effect of ApTOLL in other TLRs. TLRs activity assay in HumanTLR2, -3, -4, -5, -7, - 8 and -9 expressing cell lines. No agonistic effect was detected after incubation with ApTOLL (20 and 200 nM). **B-C:** Histograms of ApTOLL (off-target). Incubation with ApTOLL (20 nM) showed no inhibitory effect on the activation of any target selected neither GPCRs, Ion Channels, Kinases, Nuclear Receptors, Transporters nor other Non-Kinase Enzymes. Uptake results and binding assays are represented. Results of the one-way ANOVA with a Dunn’s or Tukey’s *post hoc* test are represented as: *p <0.05.

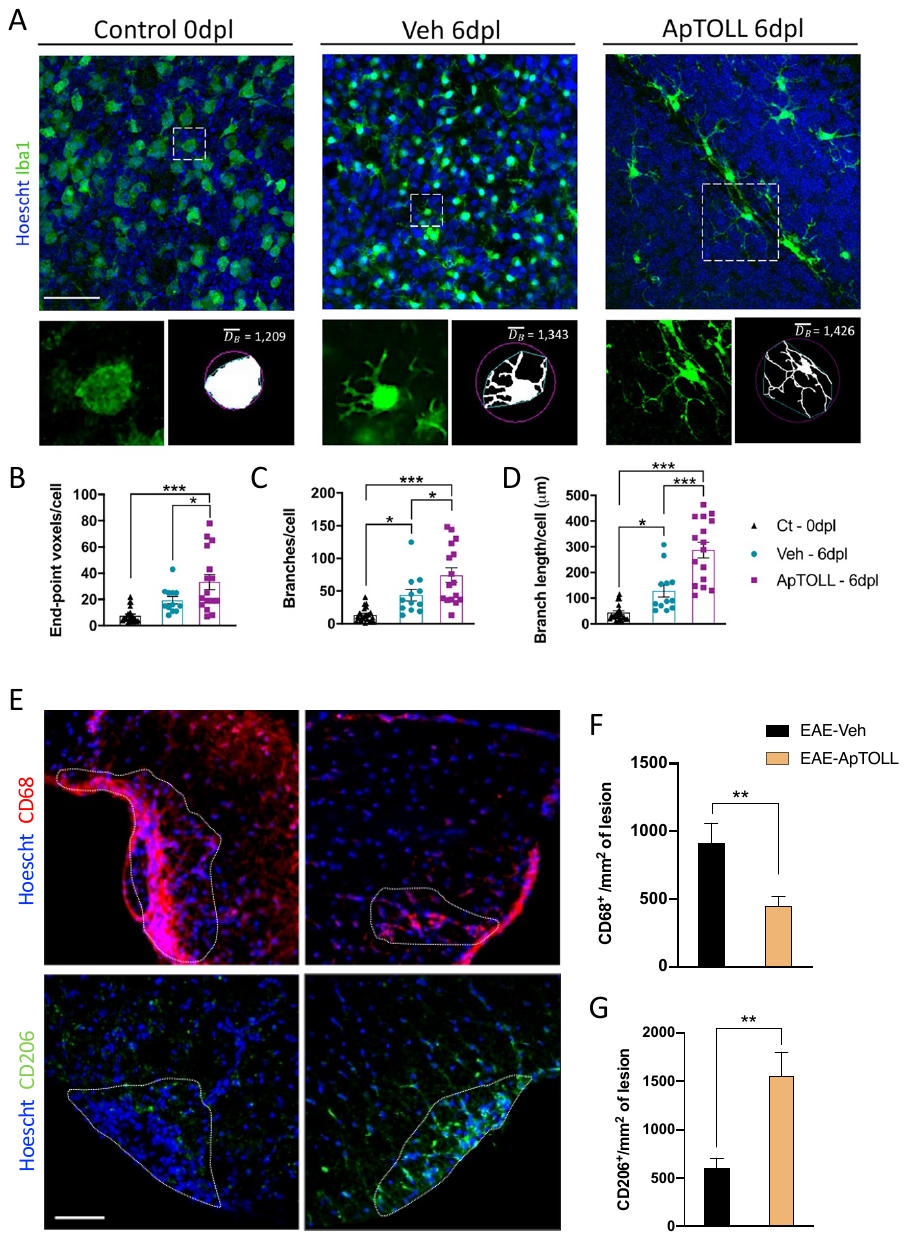

**Supplementary Fig 4. ApTOLL favors the morphological shift of Iba1^+^ cells to an anti-inflammatory phenotype**. **A:** Representative images of microglial cells and magnified images of single cells selected randomly from organotypic cerebellar slices exposed to LPC. **B-D:** Skeleton analysis of cells shows an increase in the end-point voxels (**B**), branches (**C**) and in their length (**D**) when treated with ApTOLL. The results reveal an activated state in the control condition at 0 dpl and a complex/ramified morphology in ApTOLL treated slices. At least 30 cells were analyzed in a minimum of three independent experiments.Scale bar, 50 μm in A and 15 μm in magnifications. The results of the one-way ANOVA are represented as: *p <0.05, and ***p <0.001.

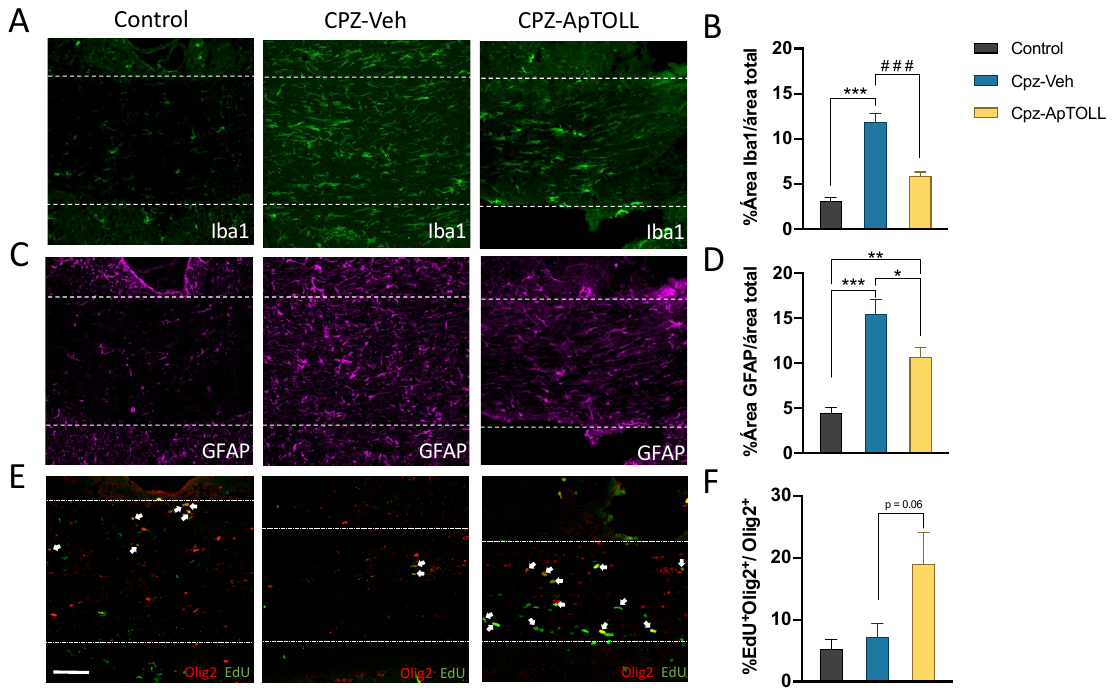

**Supplementary Fig. 5.** **ApTOLL decreases inflammatory processes and astrogliosis but increases OPCs proliferation after 3 weeks of cuprizone diet**. **A-D:** Representative images and quantification of microglia (**A-B**) and astrocytes (**C-D**) showing the percentage of the area occupied by each marker with respect to the total CC area analyzed. **E-F:** Analysis of proliferating oligodendrocytes during the third week of cuprizone. Scale bar, 60 μm. Sample size was at least 4 mice per group. Results of 1-way ANOVA with a Dunn's or Tukey's post hoc test are represented as: *p <0.05; **p <0.01, and ***p <0.001. Significant results of Student's t-test comparing pairs of groups are indicated as: ### p <0.001.

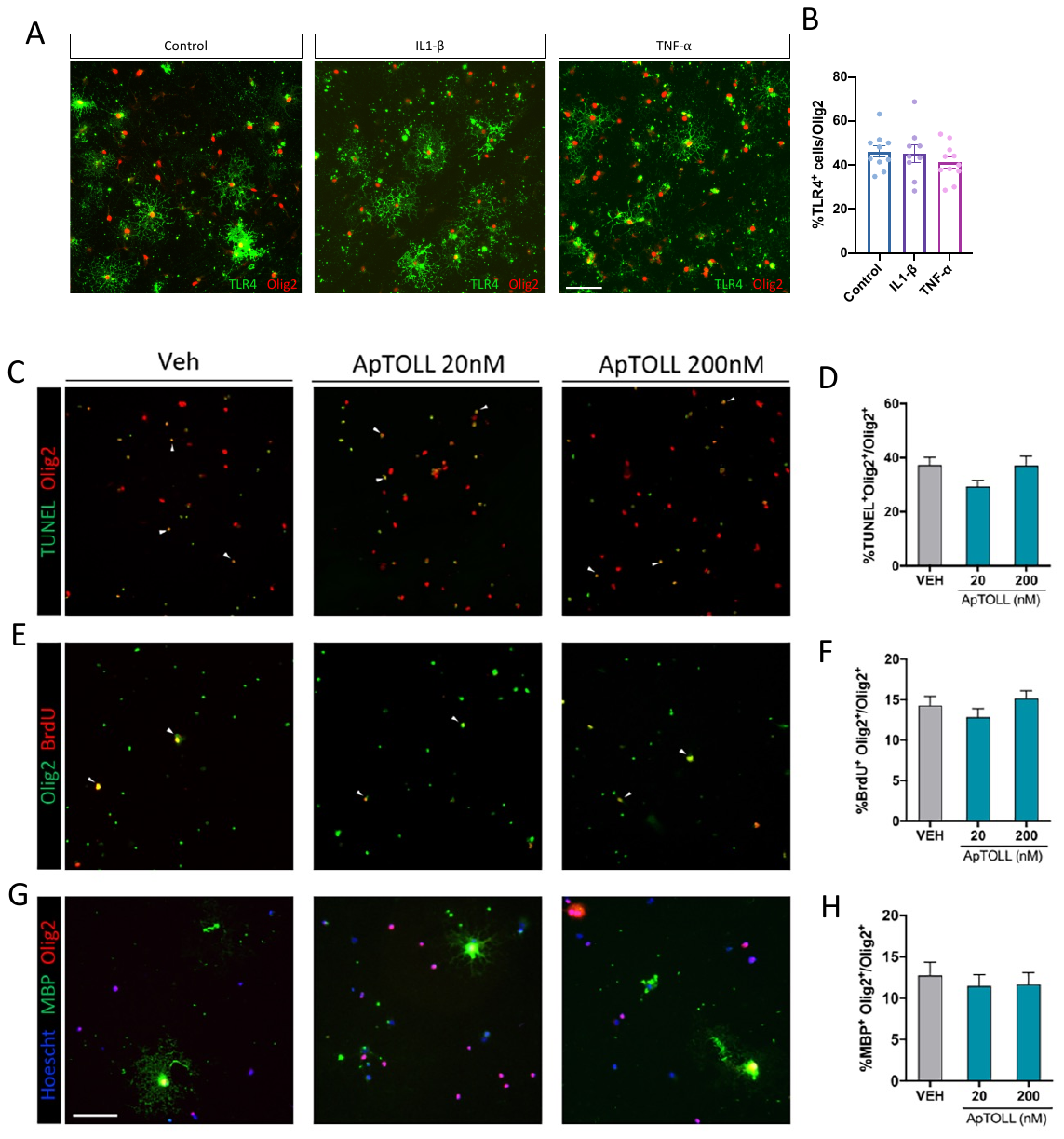

**Supplementary Fig 6. ApTOLL exerts an effect in the proliferation and differentiation of OPCs through TLR4. A-B:** Representative images of TLR4 expression in OPCs under inflammatory conditions. No differences between control and cells treated with IL1-β or TNF-α were observed. **C-H:** The OPC cultures from TLR4^-/-^ mice revealed no differences between any group regarding survival (**C, D**), proliferation (**E, F**) or differentiation (**G, H**). Scale bar, 60 μm in A, C, E and G, and one-way ANOVA was performed with a Dunn’s or Tukey’s *post hoc* test.
